# Drugs Commonly Used in Fungal Infection Risk Conditions Promote Drug Tolerance or Resistance in *Candida albicans*

**DOI:** 10.1101/2024.11.09.622705

**Authors:** Mariella Obermeier, Margy Alejandra Esparza Mora, Olivia Heese, Nir Cohen, Sreejith Jayasree Varma, Pinkus Tober-Lau, Johannes Hartl, Florian Kurth, Judith Berman, Markus Ralser

## Abstract

**Background:** Fungal infections are an increasing concern, particularly among immunocompromised patients and those with comorbidities who require multiple medications. However, the effects of drugs targeting human pathways on fungal cells, and whether they influence antifungal drug responses, are poorly understood.

**Methods:** We systematically analyzed clinical guidelines to shortlist non-antifungal drugs commonly used in conditions that increase the likelihood of fungal infections. Focusing on the most prevalent fungal pathogen, we then tested how these drugs affected the antifungal response of *Candida albicans* to two commonly used antifungals, fluconazole and anidulafungin. Drug interactions identified were further assessed using checkerboard and disk diffusion assays. Finally, antifungal treatment efficacy of fluconazole in combination with negatively interacting drugs was evaluated in an *in vivo Galleria mellonella* model of disseminated *C. albicans* infection.

**Findings:** Out of 119 drugs frequently co-administered with antifungals in 40 conditions associated with a high risk of fungal infections, 34 compounds affected the antifungal drug response in *C. albicans*, with most drugs reducing or antagonising antifungal efficacy, several through increasing resistance or tolerance. Notably, fluconazole combinations with carvedilol and loperamide promoted antifungal resistance in both fungal cultures and in *Galleria mellonella*.

**Interpretation:** Our findings suggest that medications frequently taken by patients at risk of fungal infections regularly act on the fungal pathogens and can affect the effectiveness of antifungals. We propose that human drugs acting on fungal pathogens may be an underestimated factor contributing to the evolution of antifungal tolerance and resistance.

## Introduction

Fungal infections represent a substantial medical challenge, affecting more than one billion people globally ^1^. The infections range from mild skin or nail conditions, to irritating mucosal or vaginal infections, to an increasing number of life-threatening invasive disease. Recognizing the urgent need to address the global health burden of these infections, the World Health Organization released the first Fungal Priority Pathogens List in 2022, ranking the ascomycete yeast *Candida albicans* in the highest category of critical pathogens ^2^. Indeed, *C. albicans* is a leading cause of mortality in intensive-care units, and up to 40% of the most severe invasive infections fail to respond to antifungal treatment ^3,4^. Resistance against antifungals is a known but still rare contributor to these treatment failures, while antifungal tolerance is commonly observed, but its association to treatment failure is less clear ^5^.

Some individuals are particularly at risk for a life-threatening progression of fungal infections. These include patients with immunosuppression, those who have undergone abdominal surgery, those in intensive care units, and those with severe wounds ^3,6,7^. Many of these conditions require complex medical regimens. The resulting polypharmacy can trigger drug-drug interactions (DDIs) that affect the efficacy of the antifungal drugs ^8^. For example, an antagonism between the azole antifungal fluconazole (FLC) and the antibacterial sulfadiazine increased the growth of *C. albicans* cells present in a liquid culture compared to the culture containing only FLC ^9^.

Despite the recognition that DDIs inducted by drugs directed against human targets can alter antifungal responses in the pathogen, only a limited number have studied the role of DDIs in the context of antifungal responses systematically, and these mostly focussed on the possibility of drug repurposing ^10–13^. Moreover, there is limited data available on ‘drugs of clinical relevance’ that are frequently administered to patients who suffer from fungal infection as a comorbidity or as a treatment side effect. To address these issues, we mined the medical guidelines collected by the Association of the Scientific Medical Societies (AWMF) in Germany ^14^, for references to fungal infection comorbidities alongside a wide range of disease. We used the guidelines to identify and to prioritise most commonly administered non-antifungal drugs co-prescribed to patients with fungal infections or risk of fungal infections. We then tested 119 shortlisted compounds in combination with standard antifungals in *C. albicans* cultures. We report multiple DDIs with both tested antifungals, FLC and anidulafungin (ANI). While a small number of drugs increased the efficacy of the antifungals, we obtained a higher hit rate of drugs that negatively influenced the efficacy of the antifungal. This included several interactions which enhanced drug resistance and/ or tolerance of the pathogen *in vitro* and reduced antifungal treatment efficacy in a simple *in vivo* model for invasive *candidiasis*.

## Results

### Mining health care guidelines to identify compounds most commonly co-administered clinically during fungal infection comorbidities

We systematically mined the clinical guidelines collected by the Association of the Scientific Medical Societies in Germany (Arbeitsgemeinschaft der Wissenschaftlichen Medizinischen Fachgesellschaften, AWMF (https://www.awmf.org/ ^14^) to identify pathologies or medical interventions that state a relationship with fungal infections (**Figure 1A**). In a first step, the total set of 813 clinical guidelines accessible through the AWMF repository were obtained. These guidelines were parsed computationally for the presence of 75 terms related to fungal infections or their treatment, including, e.g., “fungal infection”, “Candida” or “antifungal”. At least one keyword was detected in 249 guidelines, amounting to approximately one third of all screened guidelines.

**Figure 1:**
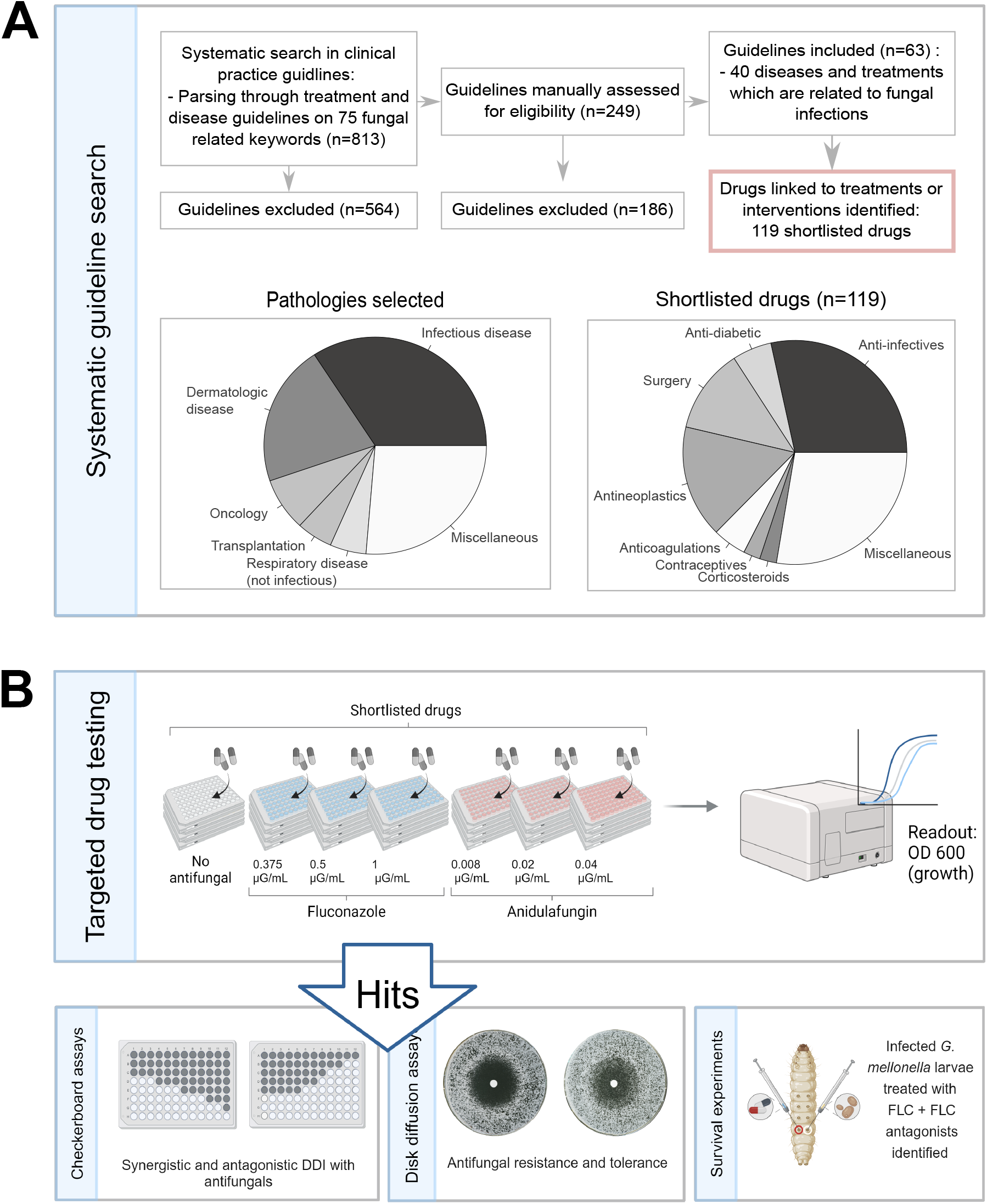
Workflow of the medical guideline search and experimental set-up. **(A)** To identify pathologies in fungal infection high risk settings, 813 medical guidelines were systematically mined on their association with fungal infections. The search resulted in 63 guidelines which indicate a relation of 40 pathologies or medical interventions to fungal infections. A total of 119 drugs which are commonly used in these identified pathologies and interventions were shortlisted. Pathology classes selected in the systematic keyword search and related shortlisted drugs are shown in pie charts. **(B)** While targeted drug testing, the shortlisted drugs were systematically exposed to *C. albicans* SC5314 cultures in presence and absence of the antifungals fluconazole (FLC) and anidulafungin (ANI). Hits, defined as compounds which increased or decreased the OD600 after 60h culturing, were further tested in checkerboard assays to identify synergistic and antagonistic DDIs, and disk diffusion assays to determine whether the interactions altered antifungal tolerance and/ or resistance. Finally, FLC antagonists identified from checkerboard assays were co-administered to FLC treatment in *G. mellonella* larvae infected with *C. albicans*. Larvae survival was tested every 24h.

In a second step, the 249 associations were manually assessed for content to exclude false positives. Guidelines were included in the final list if they addressed pathologies or medical interventions stated as increasing the risk of fungal infections, or if they could be caused by fungal infections (**Suppl. Table 1**). The severity and type of fungal infection was not considered in the selection process, but guidelines were excluded if they had no medically relevant connection to fungal infections. In total, we obtained a set of 63 guidelines addressing the treatment of 40 pathologies or medical interventions where fungal infections often occur (**Figure 1A & Suppl. Table 1**). For example, the guideline on how to treat multiple myeloma noted invasive fungal infections as a severe complication and a prognostic factor for the severity of multiple myeloma, and recommended an anti-infection prophylaxis dependent on the conducted therapy and individual risk factors (**Suppl. Table 1**, AWMF guideline registration number 018/0350L ^15^).

**Table 1:** Compounds affecting the antifungal responses of *C. albicans*.

| Compound name | Suggested MOA<sup>A</sup> | Relevant health Conditions | FLC<sup>res</sup> (24h) | FLC<sup>tol</sup> (72h) | *G. mellonella* mortality increase | Relevant citations |
| --- | --- | --- | --- | --- | --- | --- |
| **Ethinyl estradiol** | B-estradiol derivate | Contraceptive | ↑ | ↓ | **Yes**<br>($p < 0.001$) | Steroid hormones estrogen and progesterone protect *Candida* cells from FLC by increased *CDR1* gene expression<sup>31,32</sup> |
| **Desogestrel** | Synthetic gestagen | Contraceptive | → | ↑ | No |  |
| **Dienogest** | Synthetic gestagen | Contraceptive | ↑ | ↓ | Not statistically significant |  |
| **Levothyroxine sodium** | Synthetic thyroxine (T4) | Treatment hypothyroidism | → | ↓ | **Yes**<br>($p < 0.01$) | Previous drug screen also reports reduced antifungal effect of FLC on fungal growth<sup>12</sup> |
| **Loperamide hydrochloride** | Opiate receptor modulator | Chemotherapy-related diarrhoea | ↑ | ↓ | **Yes**<br>($p < 0.01$) | none |
| **Doxorubicin hydrochloride** | Topoisomerase II inhibitor | Antineoplastic agent | ↑ | ↑ | No | Increased *CDR1* and *CDR2* gene as well as *Cdr1p* and *Cdr2p* protein expression<sup>33</sup> |
| **Salmeterol xinafoate** | Adrenergic β-2 receptor agonist | COPD and asthma | ↑ | ↓ | No | Previous drug screen also reports reduced antifungal effect of FLC on fungal growth<sup>12</sup> |
| **Carvedilol** | Non-selective β-blocker | Liver cirrhosis and rosacea | ↑ | ↑ | Not statistically significant | Previous drug screen reports reduced fungal growth after 24h<sup>12</sup> |
| **Mycophenolate mofetil** | Inhibits Inosine-50-monophosphate dehydrogenase | Transplants (immune suppression) | NA<sup>B</sup> | ↓ | No | Antagonism of MPA<sup>C</sup> + FLC in *C. glabrata*<sup>34</sup> |
| **Tacrolimus** | Calcineurin inhibitor | Transplantation (immune suppression) | ↑ | ↓ | Not statistically significant | Clears tolerance in vitro, presumably via stress responses<sup>30,35</sup> |

Footnotes: A: MOA = Mechanism of action, B: NA = not sufficient cell growth after 24 h to measure the RAD, C: MPA = mycophenolic acid.

Next, we used the selected guidelines to extract the list of (non-antifungal) drugs used in these pathologies as part of standard care. We ranked the compounds based on the frequency of occurrence, and also prioritised different drug classes to cover multiple mechanisms of action. This process resulted in a shortlist of 119 (non-antifungal) drugs (**Suppl. Table 2**) which included compounds used in oncology and transplantation medicine, but also compounds that are less commonly associated with a fungal health burden disease, such as muscle relaxants or antihistamines. Furthermore, a significant proportion of the drugs reflect the common problem of fungal - bacterial, and fungal - viral co-infections (**Figure 1A**, “Infectious diseases’’), and thus, the list also included bacterial antibiotics and antivirals (n=35, **Figure 1A**, “Anti-infectives”).

### Testing the effect of clinically used compounds for interactions with antifungals

The 119 targeted compounds were obtained commercially (**Suppl. Table 2**), dissolved in DMSO or water, depending on their solubility properties. Next, we measured their effect on the growth of *C. albicans* lab strain SC5314 compared to a vehicle control, the compound administered alone (compound control) and in combination with the antifungals FLC and ANI. This setup thus allowed us to test for potential DDIs with an azole and an echinocandin drug, respectively, which are often the first-line drug classes used to treat systemic *C. albicans* infections ^16^. Drug concentrations used ranged from no drug, to below and above the MIC50 (**Figure 1B**). Fungal growth (OD_600_) was measured over 60 hours culturing and drug combinations of compound and antifungal were classified as a “hit” if the mean of all tested replicates (n=3) enhanced or reduced the biomass of the fungal cultures by 1.5- or 0.5-fold compared to the mean of the control (antifungal + vehicle) in the same growth condition.

#### Effect of targeted clinical compounds: intrinsic antifungal activities and synergism with FLC and ANI

Two of the 119 screened compounds, sirolimus and octenidine dihydrochloride, had intrinsic antifungal activity and reduced *C. albicans* growth without an added antifungal drug (**Suppl. Figure 1**). Further 22 compounds (∼18% of all tested compounds) altered *C. albicans* growth when given in combination with an antifungal drug (**Figure 2 A&B and Suppl. Figure 2**). Of these, n=12 (∼55%) affected *C. albicans* only in combination with FLC, n=1 (∼4%) were effective only in the presence of ANI, and n=9 (∼41%) affected growth in combination with both, FLC and with ANI (**Figure 2B**).

**Figure 2:**
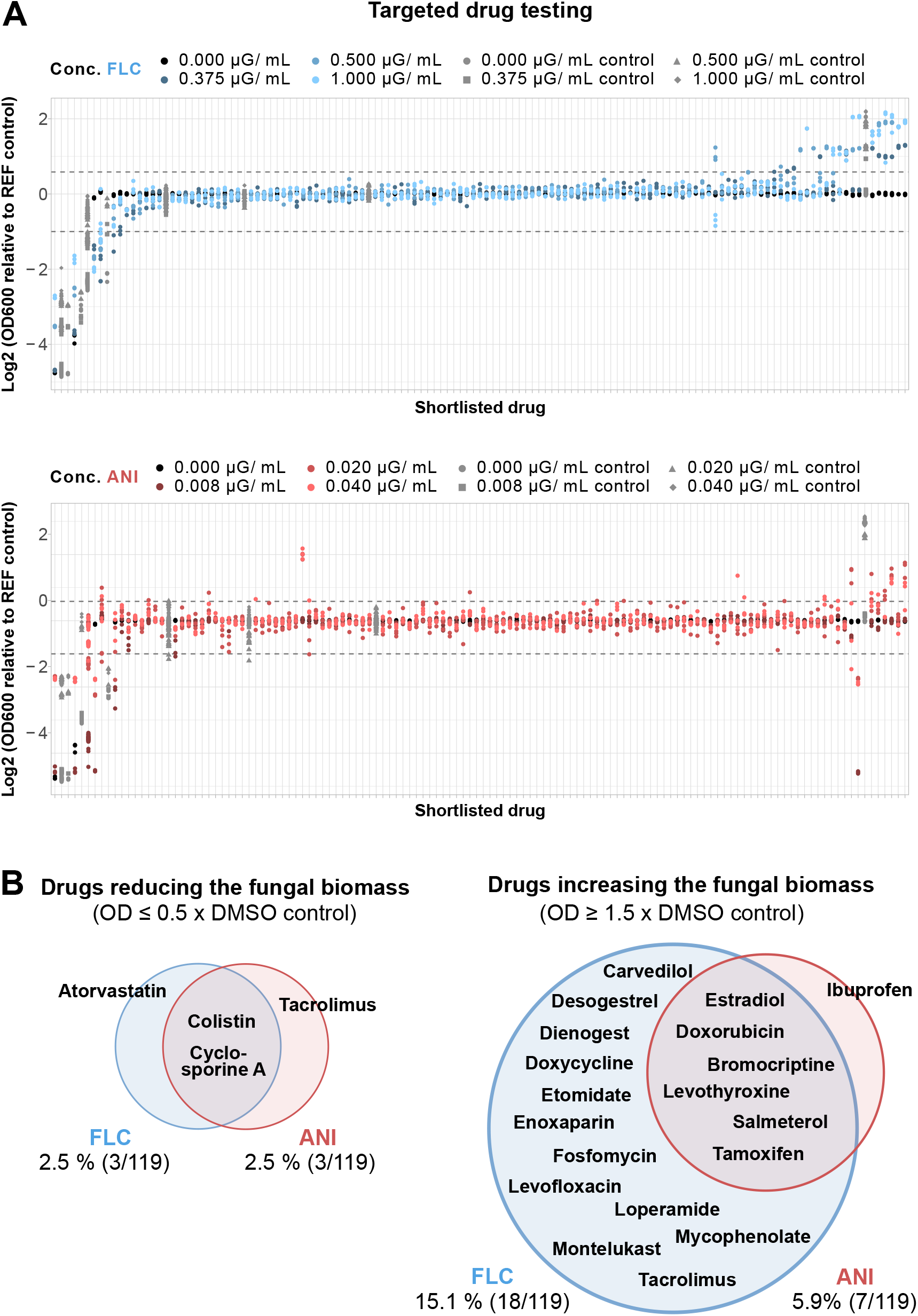
Targeted drug testing in *C. albicans* reveals negative and positive interactions of human drugs commonly administered in conditions with risk of fungal infections. **(A)** Shortlisted drugs were exposed to *C. albicans* cultures, in absence and presence of different concentrations of fluconazole (FLC, blue) and anidulafungin (ANI, red). Fungal growth, quantified by measuring the OD600 after 60 h culturing, of the tested shortlisted drugs is displayed as Log2 of the relative growth compared to a REF control, being the antifungal + DMSO of the matched condition. Grey dashed lines represent the threshold defining a “hit”. Compounds were tested in n=3 technical replicates. Detailed plots on targeted drug testing are displayed in **Suppl. Figure 2. (B)** Hits identified from targeted drug testing. Interactions with FLC (blue) and ANI (red) classified as “hits” altered the fungal OD by more than 0.5x and 1.5x mean of OD600 compared to the mean of the reference control (antifungal + DMSO).

Four compounds (∼3 % of all tested compounds) increased the antifungal activity of FLC and/ or ANI. These drugs included two immune-suppressing calcineurin inhibitors (cyclosporine A and tacrolimus ^17^), one HMG CoA reductase inhibitor (atorvastatin ^18^) and one antimicrobial that targets membrane integrity in Gram-negative bacteria (colistin sulphate ^19^, **Figure 2B**). Checkerboard assays of these positive interactions (**Figure 3A**) showed that, as seen in prior studies ^20–22^, atorvastatin and colistin sulphate interacted synergistically with FLC, and that cyclosporine A and tacrolimus interacted synergistically with ANI (**Figure 3B**). The other drug-antifungal combinations, including cyclosporine A - FLC and colistin sulphate - ANI, did not show definitive synergistic interactions (**Suppl. Figure 3a**).

**Figure 3:**
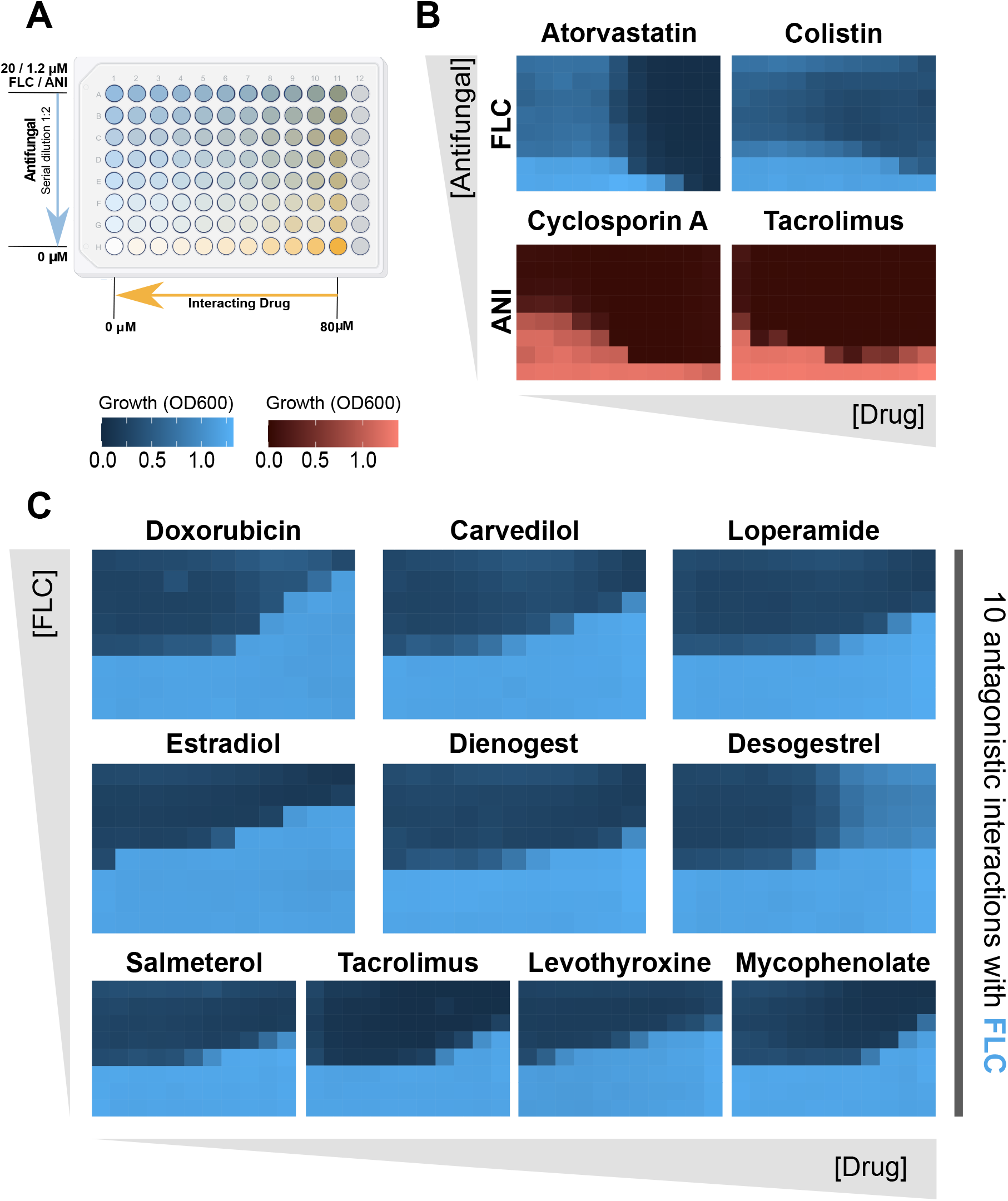
Checkerboard assays identify synergistic and antagonistic drug interactions with FLC and ANI. **(A)** Experimental set-up of checkerboard assays, including a concentration range of the antifungal (fluconazole (FLC) or anidulafungin (ANI)) on the y axis and concentration range of the interacting drug on the x axis of a 96-well edge plate. **(B)** Checkerboards with FLC (blue) and ANI (red) of drugs that suppressed the fungal biomass of at least two steps downwards the typical staircase pattern of synergistic DDIs. **(C)** Checkerboards of ten drugs that increased the biomass of *C. albicans* show the typical staircase pattern of antagonistic DDIs, being at least two steps upwards. **(B)** and **(C)** display one of the experimentally conducted two biological replicates. All performed checkerboards are shown in **Suppl. Figure 3**. All checkerboards were tested with two biological replicates.

#### Drug antagonism of ten commonly used drugs with FLC

Notably, n=19 (∼16 %) of the tested compounds decreased the efficacy of the antifungals. Of these, n=12 compounds increased *C. albicans* growth at antifungal concentrations above the MIC of only FLC, n=1 of only ANI and n=6 of both FLC and ANI (**Figure 2B**). These negative interactions included several drug classes and different mechanisms of action. For example, tamoxifen and tacrolimus fully restored fungal growth at every FLC concentration tested (**Suppl. Figure 4**). In combination with ANI, ibuprofen caused the highest restoration of fungal growth, with up to ∼70% quantified biomass of the untreated control (**Suppl. Figure 4**). Among the hits were modulators of steroid hormone synthesis, including the synthetic progesterones dienogest and desogestrel ^23^, ethinyl-estradiol (a β-estradiol derivative ^24^), and tamoxifen (a selective estrogen receptor modulator ^25^). Two additional compounds disrupt DNA and RNA synthesis in prokaryotic (the antibacterial levofloxacin ^26^) or eukaryotic (the antineoplastic doxorubicin ^27^) cells. Two other identified drugs are prominent β-receptor modulators: the non-selective β-blocker carvedilol ^28^ and the β_2_-receptor agonist salmeterol ^29^.

As for the positive interactions, we conducted checkerboard assays to distinguish antagonistic and additive interactions. Drug interactions that produced a characteristic staircase pattern and resulted in a ≥ ∼4-fold change in MIC, were defined as antagonistic. Based on these criteria, ten compounds were identified as having antagonistic interactions with FLC (**Figure 3C, Table 1**). These included the hormonal contraceptives ethinyl estradiol, desogestrel and dienogest as well as, mycophenolate mofetil and tacrolimus (immune suppressors), carvedilol (an antihypertensive), salmeterol xinafoate (a bronchospasmolytic), doxorubicin hydrochloride (an antineoplastic), loperamide hydrochloride (an antidiarrheal) and levothyroxine sodium (a synthetic thyroid hormone). None of the compounds passed the ≥ 4-fold MIC change threshold with ANI (**Suppl. Figure 3b**).

### Human drugs affect both antifungal resistance and tolerance

A commonly applied definition of drug resistance in fungal pathogens is an increase in the MIC; while tolerance is defined as the slow growth of a subpopulation of cells at drug concentrations above the MIC ^30^. To distinguish whether the DDIs we detected affected FLC resistance and/or tolerance, we performed disk diffusion assays (DDAs, **Figure 4A**). In DDAs, increased resistance results in a smaller radius (RAD) of the zone of inhibition (ZOI), and tolerance is measured as the relative fraction of growth (FoG) for areas inside the ZOI (**Figure 4B**) relative to areas outside the ZOI, such that higher FoG levels reflect higher drug tolerance.

**Figure 4:**
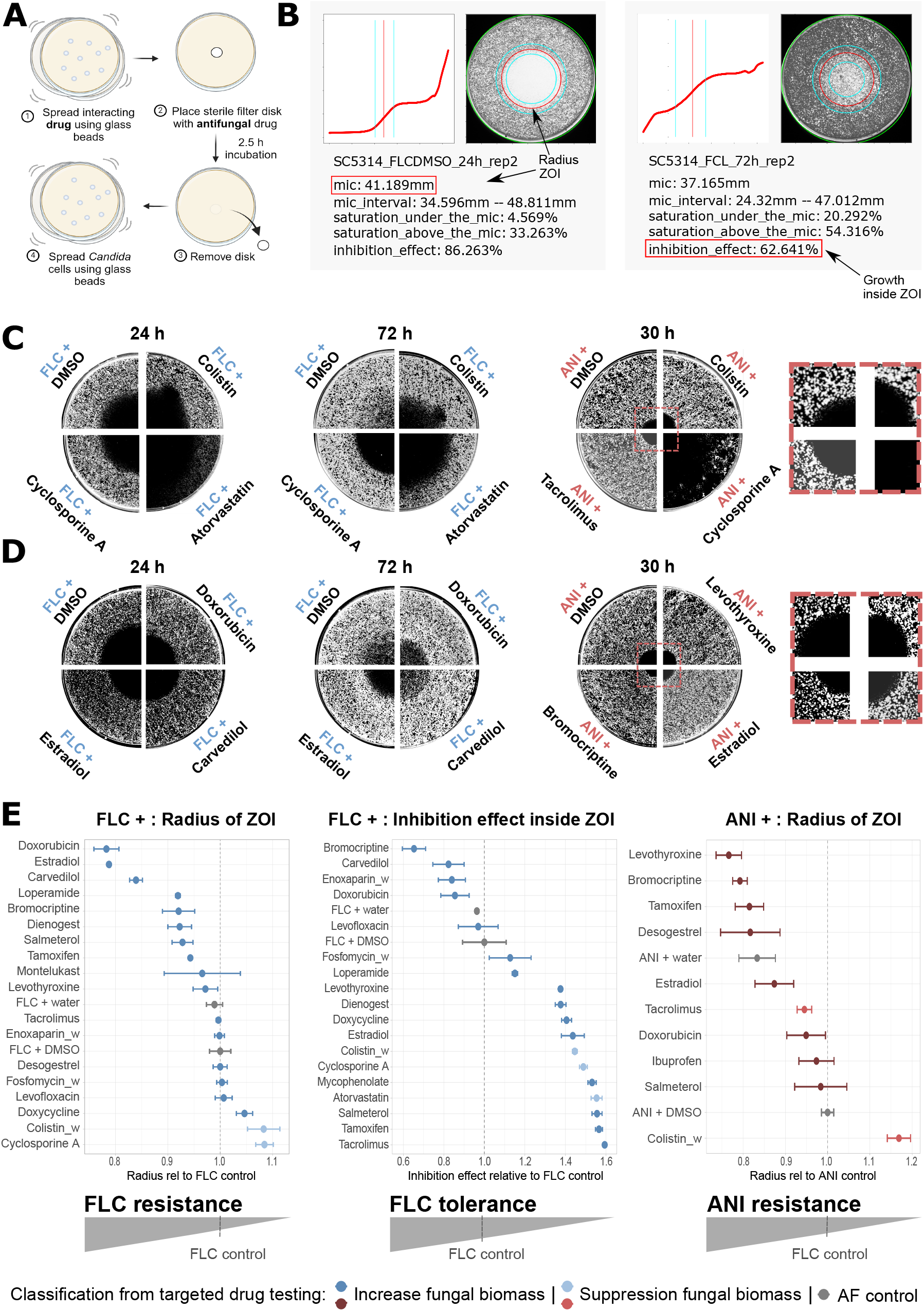
Human drugs increase antifungal resistance and tolerance in *C. albicans*. **(A)** To perform DDAs, the interacting drug was equally spread over a petri dish and subsequently, a sterile filter disk containing the tested antifungal (FLC or ANI) was placed in the center of the dish. After 2.5 h diffusion time, the disk was removed and fungal cells were equally spread over the dish. **(B)** DDAs were computationally evaluated for the radius of the ZOI after 24 h (FLC) and 30 h (ANI), corresponding to the resistance level, as well as for the inhibition effect inside the ZOI after 72h, corresponding to the tolerance level (FoG inside the ZOI). **(C)** DDAs of drugs that suppressed the fungal biomass in combination with FLC and/ or ANI during targeted drug testing. **(D)** DDAs of drugs increasing the fungal biomass in combination with FLC and/ or ANI in targeted drug tests. **(B) & (C)** All replicates (n=3 biological replicates) are shown in **Suppl. Figure 5. (E)** Many drugs altered the resistance and tolerance against FLC or ANI (antifungal + DMSO control = 1). Atorvastatin+FLC and cyclosporine A+ANI decreased FLC resistance dramatically so that a ZOI could not be detected and data are not shown here. Concomitantly, desogestrel dramatically increased FLC tolerance, so that the growth inside the halo could not be distinguished from the outer part of the ZOI, thus also not shown here (see **Suppl. Figure 5**). N=3 biological replicates of tested drugs, except DMSO+FLC (n=6). _w = drug was solved in water (instead of DMSO).

The three drugs exhibiting positive interactions with FLC (atorvastatin, colistin sulphate and cyclosporine A), resulted in increased RAD levels relative to FLC alone, and FoG levels were cleared to background levels (**Figure 4C&E**). Thus, these compounds reduced FLC resistance and tolerance levels. For drugs that positively interacted with ANI, two of three compounds (colistin sulphate and cyclosporine A) increased RAD, indicating a reduced resistance, while tacrolimus slightly reduced RAD, indicating some increase in resistance (**Figure 4C&E**). However, because the lab strain SC5314 exhibits little to no intrinsic tolerance against ANI, the potential impact of the tested drugs on ANI tolerance was not assessed.

Further, some drugs that had negative interactions with antifungals in our systematic testing were accompanied by a reduced RAD compared to the antifungal only, indicating increased resistance. In particular, bromocriptine mesylate, carvedilol, dienogest, doxorubicin hydrochloride, ethinyl estradiol, loperamide hydrochloride, montelukast sodium, salmeterol xinafoate and tamoxifen increased FLC resistance, while bromocriptine mesylate, doxorubicin hydrochloride, ethinyl estradiol, levothyroxine sodium and tamoxifen increased ANI resistance (**Figure 4D&E**). Drugs that did not show a reduced RAD, thus not indicated increased antifungal resistance, were enoxaparin sodium, doxycycline hyclate, desogestrel, fosfomycin disodium, levofloxacin hemihydrate, levothyroxine sodium and tacrolimus in combination with FLC, as well as ibuprofen and salmeterol xinafoate combined with ANI (**Figure 4E**). Interestingly, FLC tolerance assessed from drugs co-administered that caused negative DDIs with FLC showed different effects. Most drugs – including dienogest, doxycycline hyclate, ethinyl estradiol, levofloxacin hemihydrate, levothyroxine sodium, mycophenolate mofetil, salmeterol xinafoate, tacrolimus and tamoxifen – reduced or even completely removed FLC tolerance by suppressing subpopulation growth in the ZOI. Conversely, five drugs – including bromocriptine mesylate, carvedilol, desogestrel, doxorubicin hydrochloride and enoxaparin sodium – increased the FoG inside the ZOI of FLC, thus, increased FLC tolerance (**Figure 4E, Suppl. Figure 5**). Overall, negative DDIs that antagonized FLC generally also increased FLC resistance (except desogestrel), while FLC tolerance was either increased or reduced, depending on the compound.

### FLC antagonists on *C. albicans*-infected *Galleria mellonella* larvae

Next, we tested the impact of the human drugs on *C. albicans* in *G. mellonella*, chosen as a non-human/non-mammalian *in vivo* model to test whether the interaction between a human drug and the fungal pathogen has the potential to affect antifungal therapy in an *in vivo* setting. *G. mellonella* larvae were injected with *C. albicans* and subsequently treated with FLC alone or in combination with a FLC antagonist (injected drug concentration < 1mg/mL), i.e., carvedilol, dienogest, desogestrel, doxorubicin hydrochloride, ethinyl estradiol, levothyroxine sodium, loperamide hydrochloride, mycophenolate mofetil, salmeterol xinafoate or tacrolimus, respectively (**Figure 5A**). With the exception of desogestrel and mycophenolate mofetil, these drugs all limited the efficacy of the azole antifungal, and resulted in a decreased median survival time (time point when no signs of viability in 50% of treated larvae) when combined with FLC, compared to FLC alone (**Suppl. Figure 6A**). Moreover, co-administration of the drugs ethinyl estradiol, levothyroxine sodium and loperamide hydrochloride with FLC resulted in significantly higher mortality compared to those treated with FLC alone (**Figure 5B**). Neither the solvent control (PBS/DMSO) nor administration of the antagonising compound alone (Control Drug 2) affected larval survival in the absence of a fungal infection. For carvedilol, salmeterol and tacrolimus, our data revealed no statistically significant changes in survival in our experiment, but the trajectories were not identical, suggesting that drug interactions in other settings, i.e. *in vivo*, should not be ruled out, and that future studies might be warranted (**Suppl. Figure 6B**). The addition of mycophenolate mofetil, doxorubicin hydrochloride and dienogest to FLC did not impair *G. mellonella* survival at the concentrations tested. Notably, desogestrel, which antagonised FLC *in vitro*, slightly improved *G. mellonella* survival when combined with FLC relative to the FLC treatment alone (**Suppl. Figure 6B**). Thus, co-administration to FLC of three out of ten tested drugs which were antagonistic with FLC *in vitro* simultaneously significantly increased larvae mortality in a non-mammalian *in vivo* model for invasive *candidiasis*. A summarizing overview of all antagonising drugs including their effect on FLC treatment in infected *Galleria mellonella* are shown in **Table 1**.

**Figure 5:**
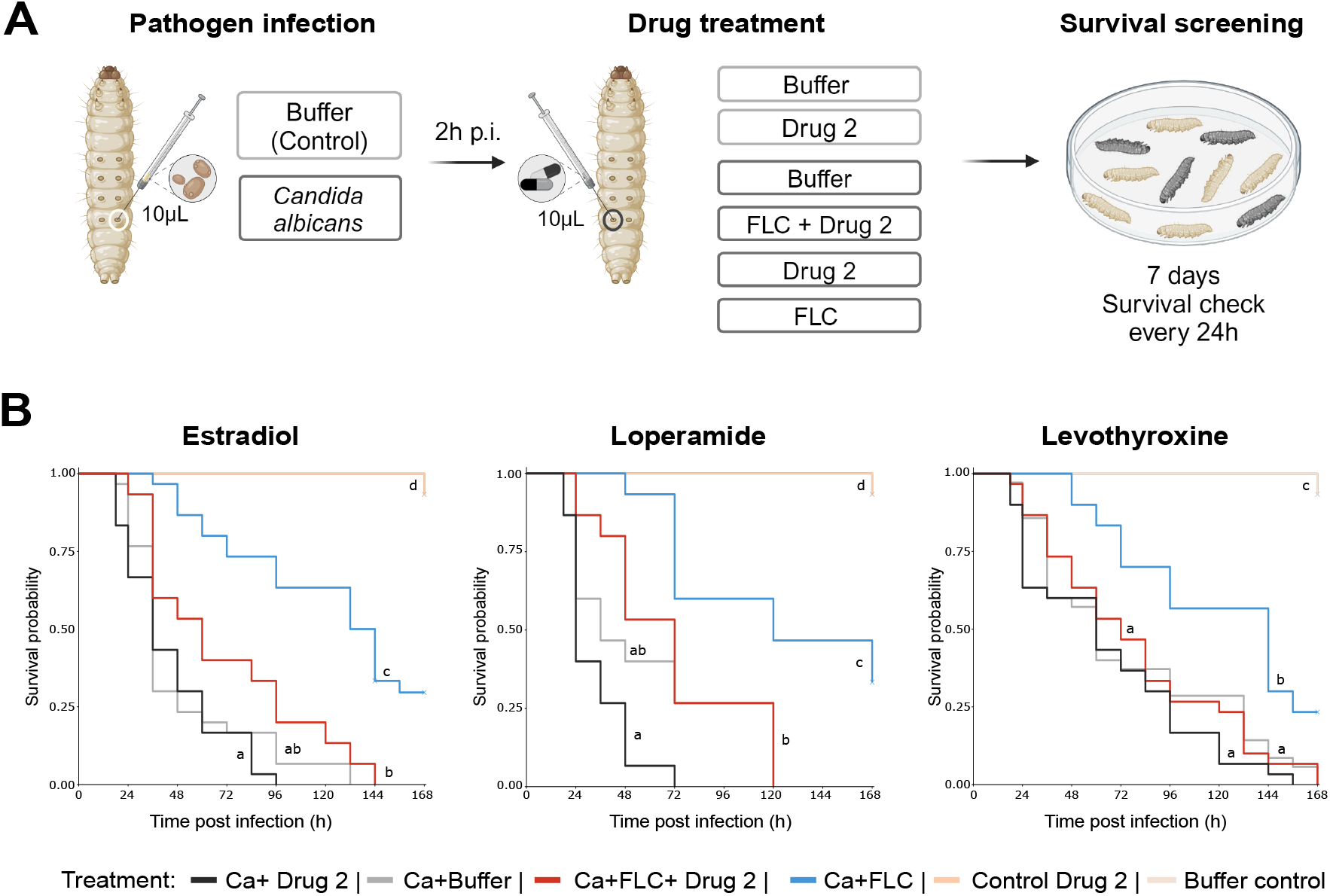
Impact of FLC antagonizers identified on *G. mellonella* survival during systemic fungal infection. **(A)** Experimental workflow of *G. mellonella* infection and drug treatment. Larvae of *G. mellonella* were injected with 5 × 10^5 *C. albicans* cells. Two hours after infection, larvae were treated with either no drug (PBS/DMSO), FLC alone or a combination of FLC + antagonising drug. Additionally, control groups of larvae were injected with PBS/DMSO and antagonizing drugs in the absence of fungal infection. After treatment, larvae were kept at 30°C and survival was monitored every 24 h. **(B)** Kaplan-Meier survival analysis of infected larvae showed significantly lower survival probabilities when infected larvae were treated with FLC and ethinyl estradiol, loperamide or levothyroxine (red) compared to FLC treatment alone (blue; Ca+FLC+Estradiol vs Ca+FLC z = 4.770 *P* = 2.76 × 10^-05, Ca+FLC+Levothyroxine vs Ca+FLC z = 3.854 *P* = 0.001746, Ca+FLC+Loperamide vs Ca+FLC z = 3.244 *P* = 0.017681). Different treatments are represented by different colours. Survival was calculated as the living proportion of larvae from every treatment group. For each condition n=15 replicates were included while each experiment was conducted twice. Crosses indicate the presence of right-censored data (i.e. larvae that were not dead at the end of the experiment). Treatment marked with the same letter showed no significant difference when performing a Wilcoxon rank sum test followed by a Bonferroni-correction to account for multiple comparisons. All statistical comparisons between each group are included in **Suppl. Table 3**.

## Discussion

While many fungal infections are mild, the risk of invasive bloodstream infections is elevated in specific patients, for instance those with comorbidities, a weakened immune system surgery or severe wounding ^3,6,7^. In these cases, treatment failures and mortality can be high. For example, mortality is ranging from 30 - 80% in *Candida* bloodstream infections ^6,7,36^. Thus far, drug resistance, a common cause of treatment failure in bacteria, is only observed in a small number (<1%) of *C. albicans* infections ^30^, but is considered to be on the rise. Meanwhile, antifungal drug tolerance, in which subpopulations can grow slowly in the presence of supra-MIC antifungal drug concentrations, is however more common, at least for fluconazole ^35^, especially at body temperature ^37^ and may underpin a proportion of antifungal treatment failures ^5,30^. Thus far we understand little, how antifungal drug resistance and tolerance evolves during severe fungal infections, and which factors influence this property.

Because fungal infections often become problematic upon medical procedures or as comorbidities in patients with poor immune status ^6^, antifungal therapies are rarely supplied alone to these patients, but rather frequently co administered on top of the primary therapies. In this study, we examined whether the coadministration of drugs used in disease with a risk of fungal infections, could affect fungal drug responses in the pathogen. Indeed, fungal and mammalian cells share many biochemical pathways ^11,38^, which creates a potential that a drug directed against a human target, has also an effect in the fungus. We focused on identifying compounds with ‘real world’ relevance, by performing a systematic search of current clinical guidelines that reference a fungal infection or an antifungal therapy. Since these guidelines are not written in a common or systematic structure, we chose a two-step approach. First, we computationally parsed the guidelines for the presence of antifungal-therapy related terms, and then we manually curated the identified guidelines for those where fungal infections indeed play a significant role. While we cannot rule out false negatives, i.e., we miss those pathologies for which the guideline does not mention fungal comorbidity, or those for which no guideline document is available, this approach offered the possibility for drug prioritisation amongst medications commonly used in routine clinical setting and in the context of fungal infections. As a result, our research mirrors the co-administration of drugs to large numbers of patients across various medical fields. Notably, this systematic strategy reflected the scale of fungal infection comorbidities; 63 guidelines of various diseases contained a reference to an increased risk of fungal infection or recommended antifungal prevention. Moreover, with this focus on negative or antagonizing interactions, our strategy was complementary to drug repurposing approaches, in some of which FDA-approved compound libraries (e.g., the Prestwick chemical library) were screened for synergism with antifungals on *C. albicans*. Of note, we define antifungal resistance relative to the control (antifungal + DMSO), rather than based on clinical breakpoints (e.g., from standardised CSLI or EUCAST guidelines ^39,40^), measured in different assays. Our study might thus detect interactions that are not seen in the standard assays applied in the clinic.

Testing 119 commonly co-administered drugs, we report an extremely high hit rate, where ∼15% and ∼6% of the drugs co-administered with FLC or ANI, respectively, affected - mostly negatively - the *C. albicans* antifungal response. Overall, FLC was more prone to be influenced by other drug treatments, an observation which is consistent with previous observations that find considerably less interactions with the tested echinocandin caspofungin compared to their observed interactions with fluconazole ^11^. For antifungal treatment, FLC is approved for treating several *Candida* infections, including vaginal and systemic infections, and is crucial for prophylaxis of *candidiasis* ^41^. A deeper understanding of the nature of these drug responses could thus be especially beneficial for optimising treatment outcomes with azole antifungals. Notably, a small number of drugs synergized with the antifungals and reduced antifungal resistance, or exhibited antifungal properties by themselves. This included compounds like sirolimus and octenidine for which antifungal activity is well documented ^21,35,42–44^, giving confidence in our approach. Indeed, sirolimus was initially discovered in a screen for antimicrobial properties measured by the diameter of the zone of inhibition in solid agar assays ^45,46^. However, although extensively discussed in literature ^47^, sirolimus has never been clinically introduced as an antifungal agent due to its immunosuppressive effects, and is currently used primarily as an immunosuppressant ^48^. Another example is colistin, which displayed synergistic interactions with FLC and ANI ^21,49^. The drug was previously shown to increase the membrane permeability of FLC-treated *C. albicans* by binding to membrane lipids. These lipids are present in the membrane of ergosterol-depleted (FLC-treated) cells ^21^.

For ten of the drugs we found antagonistic interactions with FLC, which manifested in at least two drug interaction assays: in liquid checkerboard assays, and in DDAs. Of these, doxorubicin, salmeterol and levothyroxine were previously reported to counteract FLC ^10,12^; here they increased drug resistance. To the best of our knowledge, this is the first report of antagonistic effects of FLC with mycophenolate, tacrolimus, carvedilol and loperamide in *C. albicans*.

Our shortlist included the immunosuppressants mycophenolate and tacrolimus because of their role in maintenance therapy after organ transplantation. Both drugs also weaken endogenous immune defences against infectious diseases ^50,51^. Indeed, antifungal stewardship guidelines recommend co-administration of FLC as a prophylaxis against fungal infections in certain circumstances following solid organ transplantation ^52^. Our data revealed that both drugs reduced the FLC susceptibility of *C. albicans*. Our study thus suggests future studies are warranted about the use of FLC in patients treated with mycophenolate and tacrolimus.

Loperamide, identified here as a FLC antagonist, is an opioid receptor agonist. It is frequently used to treat chemotherapy-related diarrhoea ^53^, one of the most prevalent adverse events occurring in up to 80% of patients undergoing chemo- or radiotherapy ^54^. Notably, this first-line treatment for chemotherapy-related diarrhoea is freely available over the counter in many countries and is usually consumed in extremely high doses (up to 12 and 16 mg per day for prophylactic and acute treatment, respectively) ^53^. Also in this case, our study prompts for future investigations into the use of FLC in patients receiving loperamide therapy. Another candidate highlighted by our investigations is desogestrel, a progestogenic steroid, that was the only FLC antagonist that promoted FLC tolerance rather than resistance. Here, we did not observe significant changes to viability when desogestrel was administered with FLC to *G. mellonella* larvae, but the effect on drug tolerance allows speculating that the DDI could potentially influence the evolution of drug tolerance and resistance.

Indeed, while intrinsic antifungal resistance is relatively rare in *C. albicans* strains, several of the drugs tested here promoted the acquisition of increased levels of FLC resistance as well as a few examples of increased ANI resistance. Whether the high frequency of antagonistic interactions influences the evolution of drug resistance or tolerance remains to be tested. However, it is important to highlight that the drugs tested in our study are co-administered with antifungals in millions of individuals. One concern is that many of the FLC antagonists we report - including ethinyl estradiol, desogestrel, dienogest, salmeterol, carvedilol, mycophenolate and tacrolimus - are long term medications. Prolonged usage might create conditions that increase fungal cell numbers during antifungal therapy, and thereby favour the evolution of new traits within opportunistic fungal populations. Those new traits that provide a selective advantage could then tip the scales and promote drug resistance and/or tolerance in fungal pathogens within vulnerable patients.

In summary, our study reveals a high frequency of interactions between commonly administered drugs among patients prone to fungal infections, and the prevalent fungal pathogen *Candida albicans*. Notably, many of the negative or antagonistic interactions reported involve medications that are standard treatments for conditions with a high risk of fungal infections and are frequently prescribed for long-term use in chronic diseases. The prevalence of these interactions suggests that polypharmacy may be an underestimated factor in the evolution of antimicrobial tolerance or resistance in fungal pathogens. Furthermore, our findings underscore the need for additional studies to evaluate the use of FLC in patients receiving mycophenolate, tacrolimus, or loperamide, and to determine whether these drug combinations compromise therapeutic outcomes.

## Material and Methods

### Culture conditions

All experiments were performed using *C. albicans* SC5314, a strain initially isolated from a patient diagnosed with candidiasis in the early 80s which currently serves as a global reference in laboratories doing research on *C. albicans* ^55^. Fungal cells were kept at -80 °C in 20 % glycerol YPD stocks. Two days before experimental start, *C. albicans* was streaked onto YPD agar plates and incubated at 30 °C with 60 % humidity. After 36 h of incubation, three colonies were selected and inoculated in 10 mL YPD (drug screen and checkerboard assays), followed by overnight (ON) cultivation (30 °C, 60 % humidity, shaking). Notably, in disk diffusion experiments, three colonies were individually inoculated for the ON culture. The next day, the culture was diluted 1:40 in synthetic minimal medium (SM). One litre SM was prepared from 6.7 g yeast nitrogen base without amino acids (291940, Thermo Fisher Scientific) and 2% glucose, buffered to 6.55 using 3-(N-morpholino) propane sulfonic acid (MOPS) and filled up with sterile mq H_2_O. Before starting an experiment, the pre-culture was incubated 4h (30 °C, 60 % humidity, shaking).

### Targeted drug testing

Compounds were purchased from commercial suppliers (**Suppl. Table 2**), diluted in DMSO to a working concentration of 1 mM and stored at -80 °C until usage. Compounds insoluble in DMSO were dissolved in sterile water at the day of usage to a concentration of 10 mM and diluted 1:10 in DMSO to a working concentration of 1 mM. The compound library was thawed at the day of the experiment and controls were prepared, including SM only (untreated control), DMSO only (reference control, including antifungal + DMSO), 90% DMSO / 10% water (control water soluble drugs) and supra-MIC concentration of FLC (sc-205698A, Santa Cruz Biotechnology) or ANI (SML2288, Merck). In addition, FLC, ANI and amphotericin B (sc-202462A, Santa Cruz Biotechnology) served as controls for fungal growth suppression and were used in final concentrations of 10 µM. To prepare 96-well edge plates, SM containing three concentrations of FLC (2x 0.375, 2x 0.5 and 2x 1 µg/mL), ANI (2x 0.008, 2x 0.02 and 2x 0.04 µg/mL) or an equivalent amount of DMSO (0 µg/mL FLC/ANI) were prepared and 98 µL of the antifungal / DMSO enriched medium was added to the plates. Subsequently, 2 µL of each compound from the library and corresponding controls were dispensed into the wells using the Biomek i5 robot (Beckman Coulter Life Sciences), resulting in a final concentration of 10 µM. Following a 4 h pre-culture in SM, *C. albicans* was diluted in SM to twice the inoculation concentration (OD_600_ = 2 × 0.0025). Afterwards, 100 µL of the cell solution was added followed by four times mixing in the 96-well plates. All drugs were cultured in triplicates. Additionally, an untreated (no antifungal) and blank control were incorporated in triplicate for each plate. The supra-MIC of FLC / ANI, SM only, reference and DMSO/water controls were included as quintuplicate on every plate. All conditions underwent randomization through the Python shuffle() function. Edge effects were minimised by adding 1.5 mL sterile water to every edge in the 96-well edge plate. The plates were incubated at 30 °C and 60 % humidity, and optical density at 600 nm was measured every 90 min for a duration of 72 h using a BioTek Epoch 2 microplate reader (Agilent Technologies) connected to a BioTek BioStack 3 microplate stacker (Agilent Technologies).

### Checkerboard assays

Drug interactions of the hits identified from targeted drug testing with FLC / ANI were evaluated by conducting checkerboard assays (**Figure 3A**). A 10 mM stock of the hit compound was diluted in SM to 4x 80 µM (excepting levothyroxine sodium, cyclosporine A and tacrolimus which were diluted to 4x 40 µM due to poor solubility). A 100 µL of the dissolved drug in SM was filled in a 96-well edge plate in column 11 and horizontally serial diluted (1:2) with SM, including all wells of the column 2-10. The remaining 50 µL of the drug were added to the blank positions (12A-H). Subsequently, antifungals were diluted in SM to four times the highest concentration of FLC or ANI, including 4x 20 and 4x 1.2 µM, respectively, and seven times serial diluted (1:2). Fifty µL of the dilutions were added to each well, while highest concentrations of the antifungal were added to A1-12 and systematically lowered vertically. H1-12 were filled with SM only. After a 4h pre-culture of *C. albicans*, cells were diluted in SM (OD_600_ = 2x 0.0025) and 100 µL of the cell solution was added to well A-H1-11 in the 96-well edge plates. All blank positions (A-H12) were filled up with 100 µL SM. To minimise edge effects, all edges were filled with 1.5 mL sterile water. Plates were incubated at 30 °C and 60 % humidity. OD_600_ was measured every 90 min for a duration of 72 h using a BioTek Epoch 2 microplate reader connected to the BioTek BioStak 3 microplate stacker. Two biological replicates of each checkerboard assay were prepared. Since most tested non-antifungals showed no intrinsic antifungal effect, and could thus not be assigned to a MIC, synergism and antagonism was not assessed by calculating the fractional inhibitory concentration index, as is usually the case. Instead, distinction of synergistic and antagonistic interactions from those that are additive were based on the examination of the characteristic staircase pattern of the interactions. Based on the Loewe additivity model, interactions were considered to be synergistic or antagonistic when both replicates of the assay exhibited at least two steps in the staircase pattern, roughly corresponding to an ≤ 0.5-fold (synergistic) or ≥ 4-fold (antagonistic) MIC that underlies the interpretation of the FICI ^56^.

### Disk diffusion assays

To determine the effect of identified hits on antifungal resistance and tolerance, DDAs were performed. The day before usage, 9 mm Petri dishes containing 14 mL SM-agar were prepared. A 100 µL of the hit identified during targeted drug testing at 10 mM stock concentration was evenly spread over the dish utilising 10 glass beads (4mm, 6x 10s shaking) and resulted in a final concentration of 70 µM after diffusing into the agar. Subsequently, a FLC or ANI disk containing 25 µG FLC or 5 µG ANI was placed in the center of the Petri dish. To enable diffusion of the drugs into the agar, plates were incubated for 2-3 h at RT. After incubation, the disk was removed and 100 µL of the *C. albicans* culture (OD_600_ = 0.025) were evenly distributed over the disk using 10 glass beads (4mm, 6x 10s shaking). Petri dishes were incubated for 72 h (30 °C, 60 % humidity) and scanned after 24, 30, 48 and 72 h using a Epson Perfection V800 Photo scanner (SEIKO Epson CORPORATION). DDAs were performed as biological replicates, including a blank (no fungal cells), DMSO (DMSO on the disk instead of the antifungal) and reference (FLC /ANI on the disk + DMSO evenly spread on the plate instead of the hit) control. Antifungal resistance was automatically analysed by measuring the radius of the zone of inhibition (ZOI) at 24 h (FLC DDAs) and 30 h (ANI DDAs). Antifungal tolerance was computationally evaluated in scans after 72 h by quantifying the inhibition effect inside the ZOI and correlates with colony growth within the halo (**Figure 4B**). To automate and streamline the estimation of the MIC in the DDAs, we developed a Python program that performs a radial analysis of growth patterns on the assay plate. The program applies concentric circular masks of a given thickness to summarize the gradient of growth density from the center of the assay outward. The center is determined by calculating the interpolated intersection of two perpendicular saturation curves formed by the sum over the X and Y voxel. This estimate is further refined using the signal from the diffusion disc, if present. Growth density within each circular mask is aggregated and smoothed using the Savitzky-Golay algorithm. The MIC radius is then estimated by identifying the peak of the first derivative of the resulting saturation curve, with the width of this peak corresponding to the intervals of the MIC estimation.

### *Galleria mellonella* survival infected with *C. albicans*

Antagonizing compounds with FLC identified from *in vitro* checkerboard assays were tested in an invertebrate model of *C. albicans* pathogenesis (**Figure 5A**). For this purpose, *C. albicans* was directly injected into the hemolymph of *Galleria mellonella* larvae. This procedure results in a systemic infection that enables precise testing of antifungals and other clinically relevant drugs during infection ^57,58^. Larvae were injected individually through the last right pro-leg into the hemocoel using a Hamilton Neurosyringe with either 10 µL *C. albicans* (5 × 10^7^ CFU/mL) or PBS supplemented with 10% DMSO (healthy control). Two hours post infection, larvae were treated with 10 µL of different drug combinations including FLC (4 mg/kg) + antagonizing compound: carvedilol (sc-200157, Santa Cruz Biotechnology), desogestrel (Y0000509, Sigma), dienogest (Y0001785, Sigma), doxorubicin hydrochloride (sc-200923A, Santa Cruz Biotechnology), ethinyl estradiol (sc-205318, Santa Cruz Biotechnology), levothyroxine sodium (sc-235497, Santa Cruz Biotechnology), loperamide hydrochloride (sc-203116, Santa Cruz Biotechnology), mycophenolate mofetil (sc-200971A) or tacrolimus (SEL-S5003, Biozol). FLC (4 mg/kg) and the additional compound were dissolved in PBS with 10% DMSO. Due to limited solubility of the antagonizing drugs in PBS/DMSO mixtures, the drugs were injected at saturation concentrations in 90% PBS/ 10% DMSO (injected drug concentration < 1mg/mL). One mg of each compound was sonicated in 1 mL of PBS with 10% DMSO for 10 min. Followingly, the drug-PBS/DMSO suspension was centrifuged at 14,000 rpm for 10 min and 900 µL of the supernatant was used for injection. Controls included a drug toxicity control (uninfected larvae injected with the antagonizing compound alone), a 90% PBS/ 10% DMSO control (uninfected larvae injected with the drug solvent alone), an untreated infection control (larvae infected with *C. albicans* and treated with 90% PBS/ 10% DMSO), a FLC only control (larvae infected with *C. albicans* and treated with 4 mg/kg FLC) and a drug only control (larvae infected with *C. albicans* and treated with the additional drug alone without FLC). Fifteen larvae per group were used and the experiment was repeated twice. After injection, larvae were placed in sterile 18 cm Petri dishes containing Whatman filter paper, incubated for 7 days at 30 °C and checked for survival every 24 h. Larvae that showed a brown-dark colour and no sign of movement when touched were considered to be dead. Mortality rates were calculated for each treatment. A Wilcoxon rank sum test was used to perform pairwise comparisons of mortality rate ranks between treatments and a Bonferroni-correction to account for multiple comparisons.

## Acknowledgments

This work was supported by the European Research Council (ERC) Synergy Award Fungal Tolerance (grant agreement No ERC-SyG-2020 951475 to JB and MR). P.TL., F.K. and M.R. are supported by a grant from the German Federal Ministry of Education and Research (FKZ: 031L0286A). J.B. was also supported by JPIAMR ERAnet grant CycleDrug. Figures 1, 3, 4 and 5 were partially produced using BioRender (Biorender.com).

## Data sharing

All hits of the systematic guideline search as well as the composition of the drugs shortlisted are described in Supplementary Tables. Complete results of the targeted drug testing, checkerboard assays, disk diffusion assays and *Galleria mellonella* experiments are shown in Supplementary Figures and Supplementary Tables.

## Authors contribution

M.O. contributed to conceptualization of the study and experimental design, performed the guideline search and selection of shortlisted drugs, most of the laboratory work, analysis and visualization as well as drafted the manuscript. M.R. also conceptualized the study and experimental design, supervised the study, acquired funding, and contributed to manuscript drafting, editing and reviewing. M.EM conducted the experiments, statistical analysis and visualization of all *Galleria mellonella* tests, and contributed to manuscript writing. J.B. contributed with supervision, funding acquisition and edited as well as reviewed the manuscript. O.H. performed disk diffusion assays, while N.C. developed the analysis pipeline for these assays. S.J. optimized drug preparation for the *Galleria mellonella* experiments and reviewed as well as edited the manuscript. P.TL. helped with the selection of the drug shortlist. Further, P.TL. and F.K. provided medical expertise and contributed to manuscript editing and reviewing. J.H. assisted with data visualization and manuscript editing as well as reviewing.

