## Supplementary Material for "Drugs Commonly Used in Fungal Infection Risk Conditions Promote Drug Tolerance or Resistance in *Candida albicans*"

### 1. Supplementary Tables

#### Supplementary Table 1:

Guidelines selected as “hits” after systematic keyword search in AWMF guidelines.

| AWMF guideline registration number | Guideline published/ updated | Guideline topic |
|---|---|---|
| **Guidelines including pathologies and medical interventions which indicated an association with fungal infections** | | |
| 006-052 | 2021 | Phimosis in children and adolescents |
| 007-006 | 2016 | Odontogenic infections |
| 007-096 | 2017 | Dental restoration and hear valve replacement |
| 011-021 | 2020 | Extracorporeal circulation (ECLS/ ECMO) |
| 011-022 | 2019 | Mediastinitis |
| 013-001 | 2021 | Psoriasis vulgaris |
| 013/007 | 2019 | Anal eczema |
| 013-012 | 2012 | Hidradenitis suppurative / acne inversa |
| 013-028 | 2022 | Urticaria |
| 013-033 | 2019 | Tinea Capitis |
| 013-065 | 2022 | Rosacea |
| 013-048 | 2017 | Dermatoses associated with dermal lymphostasis |
| 013-091 | 2020 | Pyoderma Gangrenosum |
| 018/0350L | 2022 | Monoclonal gammopathy, multiple myeloma |
| 018-037 | 2022 | COVID-19 in oncology and haematology |
| 020-018 | 2017 | Cystic fibrosis |
| 020-019 | 2022 | Tuberculosis |
| 021-017 | 2018 | Liver cirrhosis |
| 021-027 | 2017 | Autoimmune liver disease |
| 023-046 | 2019 | Heart transplantation |
| 026-022 | 2013 | Lung disease in cystic fibrosis patients |
| 030-044 | 2021 | HIV-1-associated neurological disorders |
| 030-050 | 2021 | MOG-IgG associated disorders |
| 030-098 | 2018 | Cerebral Thrombosis |
| 042-004 | 2016 | Visceral leishmaniasis |
| 048-014 | 2016 | Oncology |
| 057-017 | 2018 | Diabetes mellitus |
| 073-004 | 2018 | Nutrition on ICU |
| 079-001 | 2018 | Sepsis |
| 082-006 | 2017 | Parental antibacterials |
| 088-001 | 2018 | Surgery obesity and metabolic diseases |
| 112-001 | 2017 | Primary immunodeficiency |
| 113-001 | 2022 | COVID-19 |
| 151-001 | 2020 | Spondylodiscitis |

| | | |
|---|---|---|
| 166-004 | 2021 | Urinary tract infection in paediatrics |
| 189-001 | 2019 | Primary antibody deficiency |
| 018-015 | 2019 | Hormonal contraception |
| 032-054 | 2020 | Supportive treatment in oncology |
| **Guidelines including pathologies caused by fungi** | | |
| 013-048 | 2022 | Chronic Pruritus |
| 013-063 | 2020 | Pruritus ani |
| 015-071 | 2013 | Mastitis in breastfeeding population |
| 017-024 | 2015 | Tonsillitis |
| 017-049 | 2019 | Rhinosinusitis |
| 017-066 | 2019 | Antibacterial therapy in head and throat infections |
| 020-003 | 2019 | Cough |
| 020-009 | 2017 | Asthma |
| 020-013 | 2017 | Nosocomial pneumonia |
| 020-020 | 2021 | Pneumonia |
| 021-024 | 2015 | Infectious gastritis |
| 022-004 | 2015 | Infections of the central nervous system |
| 023-024 | 2014 | Infectious endocarditis |
| 024-008 | 2021 | Bacterial infections |
| 024-025 | 2015 | Neonatal sepsis |
| 027-070 | 2021 | Back pain in children and adolescents |
| 030-108 | 2021 | Brain |
| 030-0450OL | 2017 | Breast cancer |
| 043-044 | 2017 | Urinary tract infection |
| 048-011 | 2019 | HIV therapy in children and adolescents |
| 083-001 | 2018 | Urinary tract |
| 053-009 | 2014 | Earache |
| 053-010 | 2020 | Sore Throat |
| Nvl-002 | 2020 | Asthma |
| 082-005 | 2020 | *Candida* infections |

**Supplementary Table 2:**

Shortlisted drugs purchased from commercial suppliers.

| | Compound | Company | Product Number | Solvent |
|---|---|---|---|---|
| 1 | **Abacavir sulphate** | TargetMol | T6367 | DMSO |
| 2 | **Acarbose** | Santa Cruz Biotechnology | sc-203492 | DMSO |
| 3 | **Acetaminophen** | Sigma | PHR1005 | DMSO |
| 4 | **Acetylsalicylic acid** | Sigma | A5376 | DMSO |
| 5 | **Acyclovir** | Santa Cruz Biotechnology | sc-202906 | DMSO |
| 6 | **Ambroxol hydrochloride** | Santa Cruz Biotechnology | sc-200816 | DMSO |
| 7 | **Amethopterin** | Santa Cruz Biotechnology | sc-214534 | DMSO |
| 8 | **Amlodipine** | Santa Cruz Biotechnology | sc-200195 | DMSO |
| 9 | **Amoxicillin trihydrate** | Santa Cruz Biotechnology | sc-353009 | DMSO |
| 11 | **Ampicillin** | Santa Cruz Biotechnology | sc-210812 | DMSO |
| 14 | **Atorvastatin** | Sigma | PHR1422-1G | DMSO |
| 15 | **Atenolol** | Sigma | A7655-1G | DMSO |
| 16 | **Azathioprine** | Santa Cruz Biotechnology | sc-210853 | DMSO |
| 17 | **Azithromycin** | TargetMol | TGM-T6401-50MG | DMSO |
| 18 | **Baricitinib** | TargetMol | TGM-T2485-25MG | DMSO |
| 19 | **Bictegravir** | TargetMol | T4493-10mg-TM | DMSO |
| 20 | **Bromocriptine mesylate** | Santa Cruz Biotechnology | sc-200395 | DMSO |
| 22 | **Canagliflozin** | TargetMol | TGM-T1782-50MG | DMSO |
| 23 | **Capecitabine** | Santa Cruz Biotechnology | sc-205618 | DMSO |
| 24 | **Carvedilol** | Santa Cruz Biotechnology | sc-200157 | DMSO |
| 25 | **Ceftriaxone disodium salt** | Santa Cruz Bitoechnology | sc-211050 | DMSO |
| 26 | **Cefuroxime sodium** | TargetMol | TGM-T1224-100MG | DMSO |
| 27 | **Cetirizine dihydrochloride** | Sigma | 89126-50MG-F | DMSO |
| 29 | **Ciprofloxacin HCl** | Santa Cruz Biotechnology | sc-29064 | DMSO |
| 30 | **Cisatracurium besilate** | Sigma | Y0001766 | DMSO |
| 31 | **Clindamycin hydrochloride** | TargetMol | TGM-T6448-50MG | DMSO |
| 32 | **Clopidogrel sulphate** | Santa Cruz Biotechnology | sc-337638A | DMSO |
| 33 | **Cyclophosphamide monohydrate** | Sigma | C0768 | DMSO |

| | | | | |
|---|---|---|---|---|
| 34 | **Cyclosporin A** | Santa Cruz Biotechnology | sc-3503 | DMSO |
| 35 | **Dapagliflozin** | TargetMol | T2389 | DMSO |
| 36 | **Desogestrel** | Sigma | Y0000509 | DMSO |
| 37 | **Dexamethasone** | Santa Cruz Biotechnology | sc-29059 | DMSO |
| 38 | **Diclofenac sodium** | Santa Cruz Biothechnology | sc-202136 | DMSO |
| 39 | **Dienogest** | Sigma | Y0001785 | DMSO |
| 40 | **Dobutamine hydrochloride** | Sigma | D2954000 | DMSO |
| 41 | **Docetaxel** | Biomol | CDX-D0371-M025 | DMSO |
| 42 | **Dolutegravir sodium** | TargetMol | TGM-T2329-25MG | DMSO |
| 43 | **Domperidone** | Santa Cruz Biotechnology | sc-203032 | DMSO |
| 44 | **Doxorubicin hydrochloride** | Santa Cruz Biotechnology | sc-200923A | DMSO |
| 45 | **Doxycycline hyclate** | Santa Cruz Biotechnology | sc-204734B | DMSO |
| 46 | **Empagliflozin** | Santa Cruz Biotechnology | sc-482194 | DMSO |
| 47 | **Emtricitabine** | Santa Cruz Biotechnology | sc-207617 | DMSO |
| 48 | **Epinephrine** | Sigma | E4250 | DMSO |
| 49 | **Ethinyl estradiol** | Santa Cruz Biotechnology | sc-205318 | DMSO |
| 50 | **Etomidate** | TargetMol | TGM-T1089-25MG | DMSO |
| 52 | **5 - Fluorouracil** | Santa Cruz Biotechnology | sc-29060 | DMSO |
| 55 | **Furosemide** | Santa Cruz Biotechnology | sc-203961 | DMSO |
| 56 | **Ganciclovir** | Thermo Fisher Scientific | 461710010 | DMSO |
| 57 | **Glyburide** | Santa Cruz Biotechnology | sc-200982 | DMSO |
| 58 | **Hydrocortisone** | Santa Cruz Biotechnology | sc-300810 | DMSO |
| 59 | **Ibuprofen** | Sigma | PHR1004 | DMSO |
| 60 | **Imatinib mesylate** | Santa Cruz Biotechnology | sc-202180 | DMSO |
| 61 | **Isoniazid** | TargetMol | TGM-T0972-50MG | DMSO |
| 62 | **Isoflurane** | Santa Cruz Biotechnology | sc-470926 | DMSO |
| 64 | **Ketamine hydrochloride** | Sigma | K2753 | DMSO |
| 65 | **Lamivudine** | Santa Cruz Biotechnology | sc-221830A | DMSO |
| 66 | **Levofloxacin hemihydrate** | Santa Cruz Biotechnology | sc-211735 | DMSO |
| 67 | **Levothyroxine sodium** | Santa Cruz Biotechnology | sc-235497 | DMSO |
| 68 | **Lisinopril** | Santa Cruz Biotechnology | sc-205378 | DMSO |
| 69 | **Loperamide hydrochloride** | Santa Cruz Biotechnology | sc-203116 | DMSO |

| | | | | |
|---|---|---|---|---|
| 70 | **Losartan carboxylic acid** | TargetMol | T3461-25mg-TM | DMSO |
| 71 | **Melphalan** | Santa Cruz Biotechnology | sc-204799 | DMSO |
| 72 | **Metamizole sodium** | Sigma | PHR1800 | DMSO |
| 73 | **Metoclopramid** | Sigma | 32473 | DMSO |
| 74 | **Metformin hydrochloride** | Sigma | PHR1084 | DMSO |
| 75 | **Metoprolol succinate** | Target Mol | TGM-T3165-50MG | DMSO |
| 76 | **Molnupiravir** | Biozol | CMS-CS-0114880 | DMSO |
| 77 | **Montelukast sodium** | Santa Cruz Biotechnology | sc-202231A | DMSO |
| 78 | **Moxifloxacin hydrochloride** | TargetMol | TGM-T0331-50MG | DMSO |
| 79 | **Mycophenolate mofetil** | Santa Cruz Biotechnology | sc-200971A | DMSO |
| 80 | **N-Acetyl-L-Cysteine** | Santa Cruz Biotechnology | sc-202232 | DMSO |
| 81 | **Nateglinide** | Santa Cruz Biotechnology | sc-394067A | DMSO |
| 82 | **Nirmatrelvir** | TargetMol | T9351, CAS: 2628280-40-8 | DMSO |
| 83 | **Octenidine dihydrochloride** | Santa Cruz Biotechnology | sc-478739 | DMSO |
| 84 | **Omeprazole** | Sigma | 19329-50MG | DMSO |
| 85 | **Oseltamivir phosphate** | Sigma | SML1606 | DMSO |
| 86 | **Oxymetazoline hydrochloride** | Merck | O2378 | DMSO |
| 87 | **Paclitaxel** | Biomol | AG-CN2-0045-M025 | DMSO |
| 88 | **Pantoprazole sodium** | Sigma | PHR1604 | DMSO |
| 89 | **Penicillin sodium salt** | Santa Cruz Biotechnology | sc-257971B | DMSO |
| 90 | **Piperacillin sodium** | Santa Cruz Biotechnology | sc-205808 | DMSO |
| 91 | **Prednisolone** | Santa Cruz Biotechnology | sc-205815 | DMSO |
| 92 | **Propofol** | Sigma | Y0000016 | DMSO |
| 93 | **Propranolol** | Santa Cruz Biotechnology | sc-3580 | DMSO |
| 94 | **Pyrazinamide** | Santa Cruz Biotechnology | sc-205824A | DMSO |
| 95 | **Ramiprilat** | Santa Cruz Biotechnology | sc-212767C | DMSO |
| 96 | **Sirolimus** | Biomol | LC-R-5000_50mg | DMSO |
| 97 | **Remdesivir** | TargetMol | T7222 | DMSO |
| 99 | **Rifampicin** | Merck | R3501-250MG | DMSO |
| 100 | **Ritonavir** | TargetMol | TGM-T1525-50MG | DMSO |
| 101 | **Rivaroxaban** | TargetMol | T1184-25mg-TM | DMSO |
| 102 | **Rocuronium bromide** | Sigma | Y0001766 | DMSO |
| 103 | **Salmeterol xinafoate** | Santa Cruz Biotechnology | sc-202231A | DMSO |

| | | | | |
|---|---|---|---|---|
| 104 | **Sevoflurane** | Biomol (Cayman Chemical) | Cay23996-5 | DMSO |
| 105 | **SN38** | Santa Cruz Biotechnology | sc-203697 | DMSO |
| 106 | **Suxamethonium chloride** | Sigma | S2200000 | DMSO |
| 107 | **Tacrolimus** | Biozol (SellekChem) | SEL-S5003-50MG | DMSO |
| 108 | **Tamoxifen** | Sigma | 85256 | DMSO |
| 109 | **TAS102 (Trifluridine/ Tipiracil Hydrochloride)** | TargetMol | CAS 733030-01-8 | DMSO |
| 110 | **Tazobactam sodium** | Santa Cruz Biotechnology | sc-205853 | DMSO |
| 111 | **Tenofovir** | TargetMol | T1649 | DMSO |
| 112 | **Vancomycin hydrochloride** | Santa Cruz Biotechnology | sc-204938 | DMSO |
| 113 | **Warfarin** | Sigma | 45706 | DMSO |
| 114 | **Cephalexin monohydrate** | Santa Cruz Biotechnology | sc-487556 | DMSO |
| 115 | **Norepinephrine** | Merck | A7257 | DMSO |
| 116 | **Dabigatran** | TargetMol | TGM-T6295-25MG | DMSO |
| 117 | **Clavulanic acid potassium salt** | Santa Cruz Biotechnology | sc-207446 | DMSO |
| W1 | **Carboplatin** | Santa Cruz Biotechnology | sc-202093 | Water |
| W2 | **Cisplatin** | Santa Cruz Biotechnology | sc-200896 | Water |
| W3 | **Colistin sulphate salt** | Merck | C4461-100MG | Water |
| W4 | **Enoxaparin sodium** | Sigma | E0180000 | Water |
| W5 | **Fosfomycin disodium** | Santa Cruz Biotechnology | sc-211542 | Water |
| W6 | **Gabapentin** | Sigma | PHR1049 | Water |
| W7 | **Heparin sodium** | Biozol (SellekChem) | SEL-S1346-200MG | Water |
| W8 | **Oxaliplatin** | Biomol | AG-CR1-3592-M025 | Water |
| W9 | **Salbutamol hemisulfate** | Santa Cruz Biotechnology | sc-203373 | Water |
| W10 | **Tobramycin** | Santa Cruz Biotechnology | sc-204917A | Water |
| W11 | **Tranexamic acid** | Santa Cruz Biotechnology | sc-204921 | Water |
| W12 | **Tramadol hydrochloride** | Sigma | 42965 | Water |

**Supplementary Table 3:**

Statistical evaluation of *in vivo* experiments in *C. albicans* infected *G. mellonella.* ns = not significant, *p≤0.05, **p≤0.01, ***p≤0.001.

| Compound | | Calb + Drug 2 | Calb + Buffer | Calb + FLC + Drug 2 | Calb + FLC | Control Drug 2 | Buffer control |
|---|---|---|---|---|---|---|---|
| Ethinyl Estradiol | Ca + Drug 2 | - | | | | | |
| | Ca + Buffer | ns | - | | | | |
| | Ca + FLC + Drug 2 | * | ns | - | | | |
| | Ca + FLC | *** | *** | *** | - | | |
| | Control Drug 2 | *** | *** | *** | ns | - | |
| | Buffer control | *** | *** | *** | *** | ns | - |
| Loperamide hydrochloride | Ca + Drug 2 | - | | | | | |
| | Ca + Buffer | ns | - | | | | |
| | Ca + FLC + Drug 2 | ** | * | - | | | |
| | Ca + FLC | *** | *** | ** | - | | |
| | Control Drug 2 | *** | *** | *** | * | - | |
| | Buffer control | *** | *** | *** | * | ns | - |
| Levothyroxine sodium | Ca + Drug 2 | - | | | | | |
| | Ca + Buffer | ns | - | | | | |
| | Ca + FLC + Drug 2 | ns | ns | - | | | |
| | Ca + FLC | *** | ** | ** | - | | |
| | Control Drug 2 | *** | ** | ** | *** | - | |
| | Buffer control | *** | *** | *** | *** | ns | - |
| Tacrolimus | Ca + Drug 2 | - | | | | | |

| | | | | | | | |
|---|---|---|---|---|---|---|---|
| | Ca + Buffer | ns | - | | | | |
| | Ca + FLC + Drug 2 | *** | *** | - | | | |
| | Ca + FLC | *** | *** | ns | - | | |
| | Control Drug 2 | *** | *** | * | ns | - | |
| | Buffer control | *** | *** | ** | * | ns | - |
| Carvedilol | Ca + Drug 2 | - | | | | | |
| | Ca + Buffer | ns | - | | | | |
| | Ca + FLC + Drug 2 | *** | ** | - | | | |
| | Ca + FLC | *** | *** | ns | - | | |
| | Control Drug 2 | *** | *** | * | ns | - | |
| | Buffer control | *** | *** | *** | ** | ns | - |
| Mycophenolate | Ca + Drug 2 | - | | | | | |
| | Ca + Buffer | ns | - | | | | |
| | Ca + FLC + Drug 2 | *** | ns | - | | | |
| | Ca + FLC | *** | ns | ns | - | | |
| | Control Drug 2 | *** | ** | ns | ns | - | |
| | Buffer control | *** | ** | ns | ns | ns | - |
| Desogestrel | Ca + Drug 2 | - | | | | | |
| | Ca + Buffer | ns | - | | | | |
| | Ca + FLC + Drug 2 | *** | *** | - | | | |
| | Ca + FLC | *** | *** | ns | - | - | |

| | | | | | | | |
|---|---|---|---|---|---|---|---|
| | Buffer control | *** | *** | ns | * | - | - |
| Salmeterol | Ca + Drug 2 | - | | | | | |
| | Ca + Buffer | ns | - | | | | |
| | Ca + FLC + Drug 2 | *** | ** | - | | | |
| | Ca + FLC | *** | *** | ns | - | | |
| | Control Drug 2 | *** | *** | * | ns | - | |
| | Buffer control | *** | *** | ** | ** | ns | - |
| Doxorubicin | Ca + Drug 2 | - | | | | | |
| | Ca + Buffer | ns | - | | | | |
| | Ca + FLC + Drug 2 | *** | ns | - | | | |
| | Ca + FLC | *** | ** | ns | - | | |
| | Control Drug 2 | ** | * | ns | ns | - | |
| | Buffer control | *** | *** | ** | ** | ns | - |
| Dienogest | Ca + Drug 2 | - | | | | | |
| | Ca + Buffer | ns | - | | | | |
| | Ca + FLC + Drug 2 | *** | *** | - | | | |
| | Ca + FLC | *** | *** | ns | - | | |
| | Control Drug 2 | *** | *** | * | ns | - | |
| | Buffer control | *** | *** | *** | *** | ns | - |

### 2. Supplementary Figures

**Supplementary Figure 1:**

Growth rates of drugs that showed an effect on fungal growth in the presence of FLC or ANI, here exposed to *C. albicans* without any added antifungal (compound control). Data show a strong reduction in growth rate with both compounds that had intrinsic antifungal properties, including octenidine and sirolimus, but no impact on other hits. Growth rates were calculated based on the R package "growthcurver" and normalized to the average of the DMSO control of each plate (*C. albicans* growth only). N=3 technical replicates. _w = drug was solved in water (instead of DMSO).

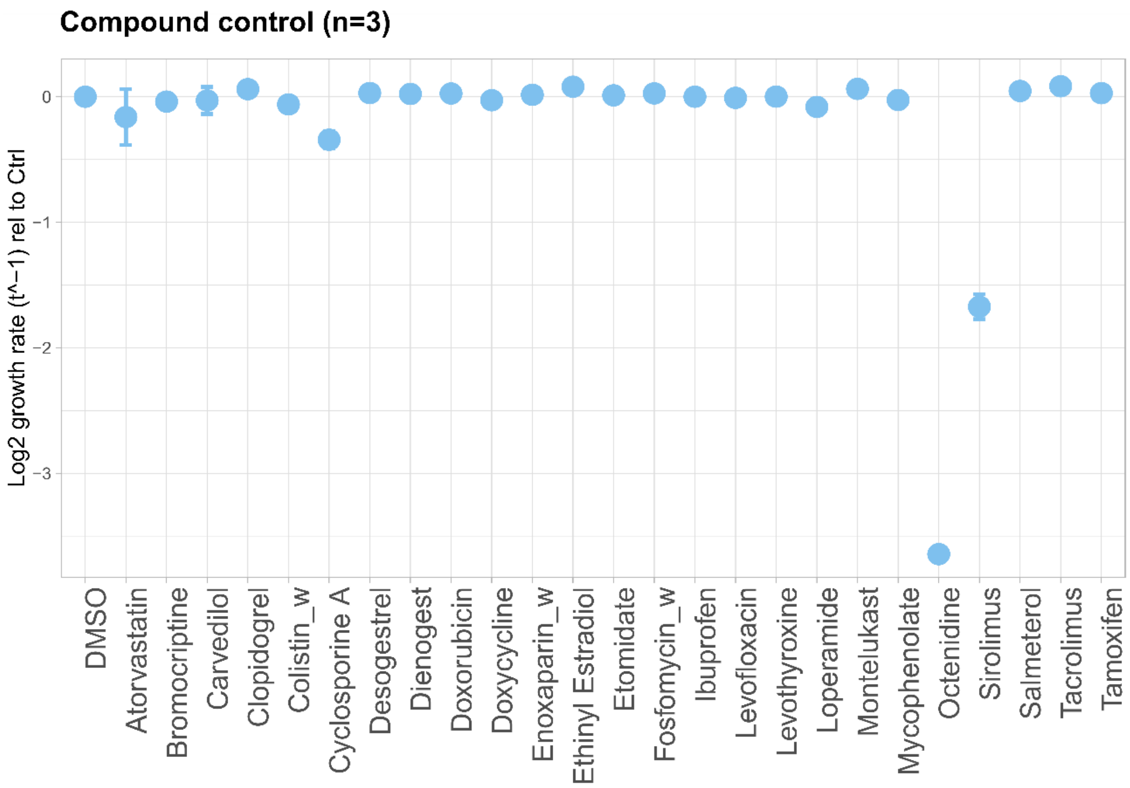


**Supplementary Figure 2**

Targeted drug testing with FLC **(A)** and ANI **(B)**. To provide a dynamic range of drug concentrations that would detect both improvement and attenuation of the FLC and ANI activities, we tested various concentrations of antifungals at i) below MIC50 (0.375 µG/ mL FLC and 0.008 µG/ mL ANI), ii) beyond MIC50 (0.5 µG/ mL and 1.0 µG/ mL FLC as well as 0.02 µG/ mL and 0.04 µG/ mL ANI) concentrations and iii) without an antifungal as a control for compounds potentially acting alone on *C. albicans*. The grey dashed line shows the threshold for a concerned "hit", having a mean of the fungal biomass > 1.5x or <0.5x OD600 of the mean of the REF control (antifungal + DMSO of the matched condition. Controls are shown in grey.

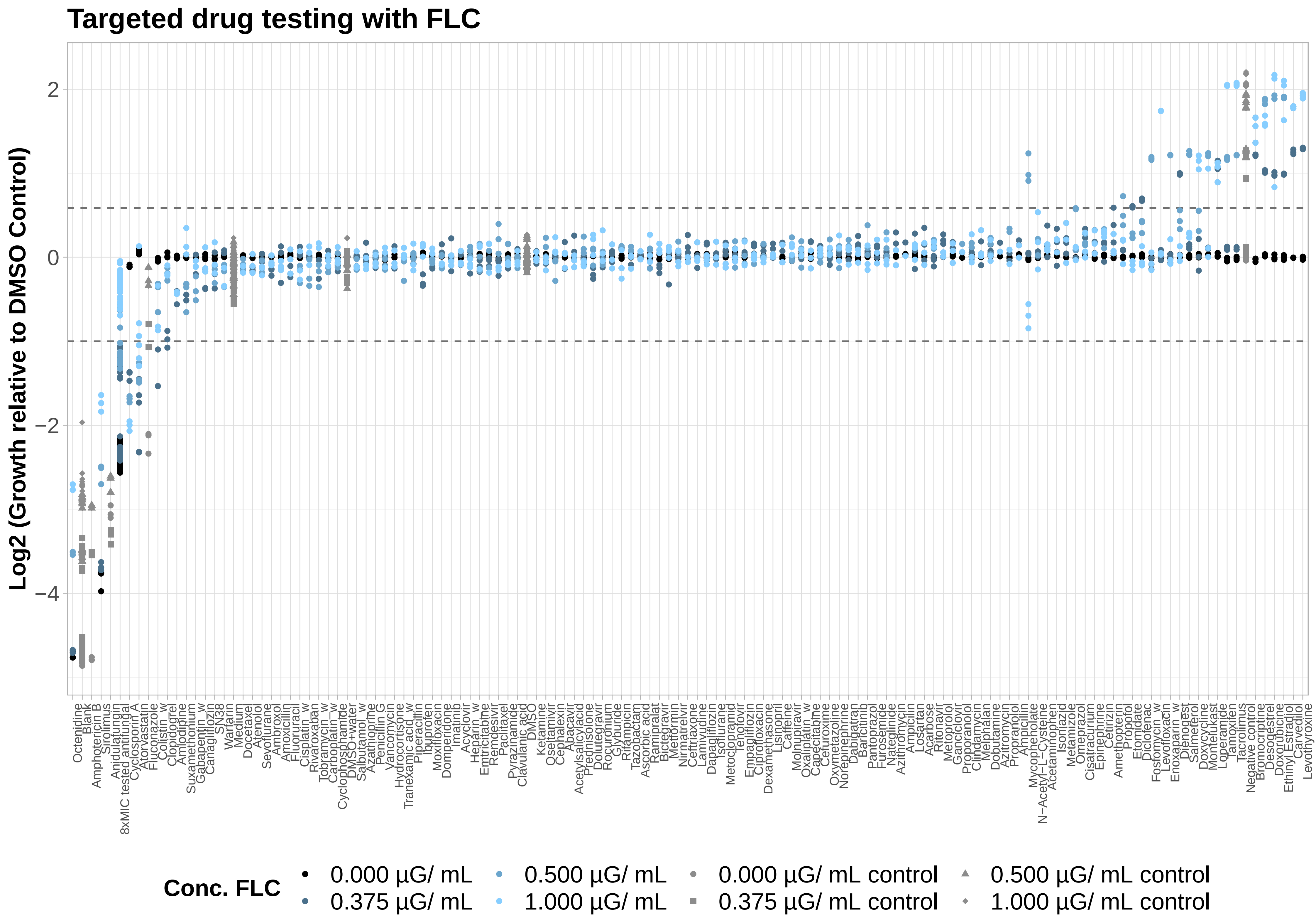

Targeted drug testing with FLC
Log2 (Growth relative to DMSO Control)
2
0
−2
−4
Octenidine
Blank
Amphotericin B
Sirolimus
Anidulafungin
8xMIC tested antifungal
Cyclosporin A
Atorvastatin
Fluconazole
Colistin_w
Clopidogrel
Amlodipine
Suxamethonium
Gabapentin_w
Canagliflozin
SN38
Warfarin
Medium
Docetaxel
Atenolol
Sevoflurane
Ambroxol
Amoxicillin
Fluoruracil
Cisplatin_w
Rivaroxaban
Tobramycin_w
Carboplatin_w
Cyclophosphamide
DMSO+water
Salbutamol_w
Azathioprine
Penicillin G
Vancomycin
Hydrocortisone
Tranexamic acid_w
Piperacillin
Ibuprofen
Moxifloxacin
Domperidone
Imatinib
Acyclovir
Heparin_w
Emtricitabine
Remdesivir
Paclitaxel
Pyrazinamide
Clavulanic acid
DMSO
Ketamine
Oseltamivir
Cephalexin
Abacavir
Acetylsalicylacid
Prednisolone
Dolutegravir
Rocuronium
Glyburide
Rifampicin
Tazobactam
Ascorbic acid
Ramipralat
Bictegravir
Metformin
Nirmatrelvir
Ceftriaxone
Lamivudine
Dapagliflozin
Isoflurane
Metoclopramid
Tenofovir
Empagliflozin
Ciprofloxacin
Dexamethasone
Lisinopril
Caffeine
Molnupiravir
Oxaliplatin_w
Capecitabine
Cefuroxime
Oxymetazoline
Norepinephrine
Dabigatran
Baricitinib
Pantoprazol
Furosemide
Nateglinide
Azithromycin
Ampicilin
Losartan
Acarbose
Ritonavir
Metoprolol
Ganciclovir
Proproanolol
Clindamycin
Melphalan
Dobutamine
Azitromycin
Propranolol
Ampicillin
Mycophenolate
N−Acetyl−L−Cysteine
Acetaminophen
Isoniazid
Metamizole
Omeprazol
Cisatracurium
Epinephrine
Cetirizin
Amethopterin
Propofol
Etomidate
Diclofenac
Fosfomycin_w
Levofloxacin
Enoxaparin_w
Dienogest
Salmeterol
Doxycycline
Montelukast
Loperamide
Tamoxifen
Tacrolimus
Negative control
Bromocriptine
Desogestrel
Doxorubicine
Ethinyl Estradiol
Carvedilol
Levothyroxine
Conc. FLC
0.000 µG/ mL
0.375 µG/ mL
0.500 µG/ mL
1.000 µG/ mL
0.000 µG/ mL control
0.375 µG/ mL control
0.500 µG/ mL control
1.000 µG/ mL control

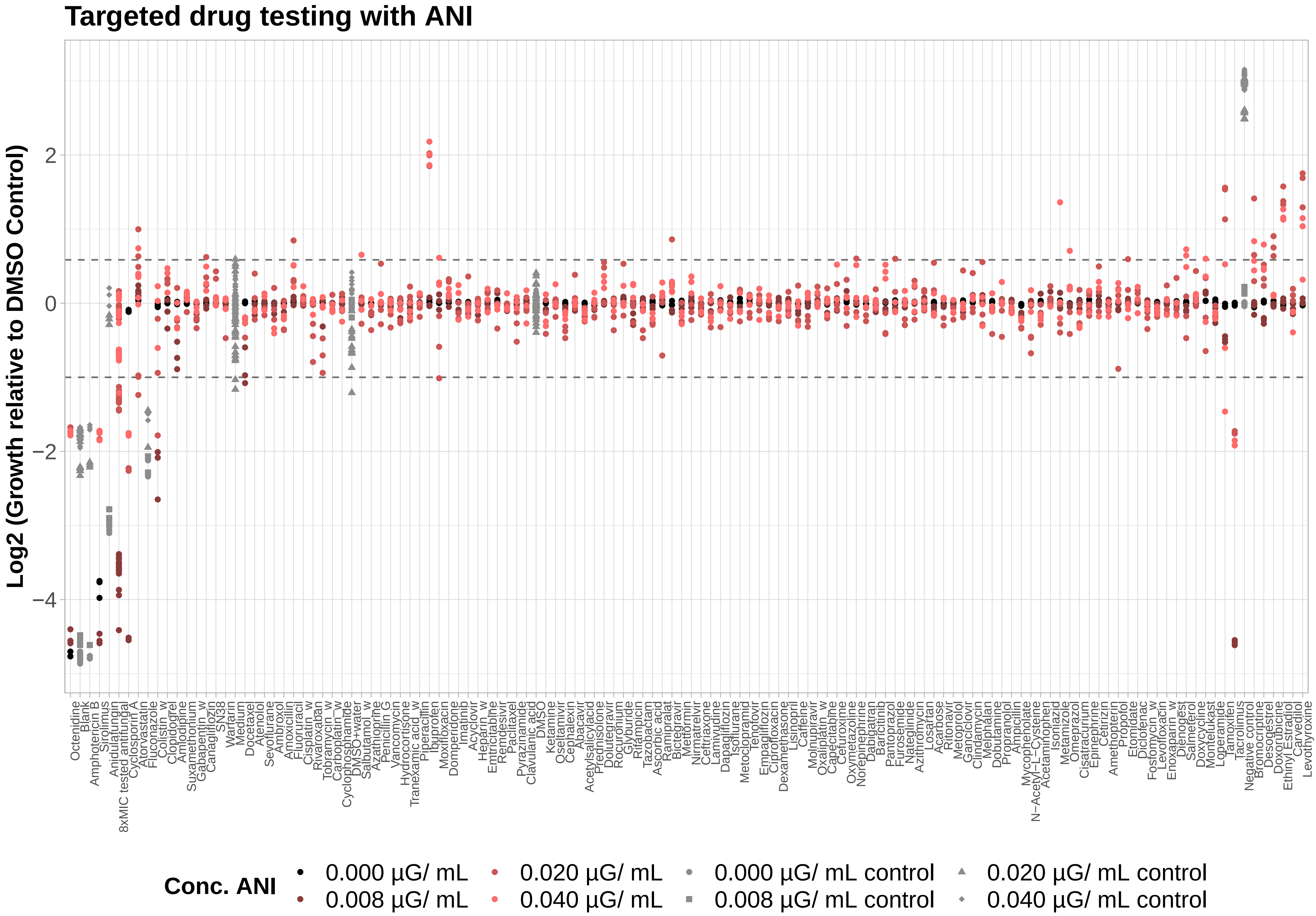

Targeted drug testing with ANI
Log2 (Growth relative to DMSO Control)
2
0
−2
−4
Octenidine
Blank
Amphotericin B
Sirolimus
8xMIC tested antifungal
Anidulafungin
Cyclosporin A
Atorvastatin
Fluconazole
Colistin_w
Clopidogrel
Amlodipine
Suxamethonium
Gabapentin_w
Canagliflozin
SN38
Warfarin
Medium
Docetaxel
Atenolol
Sevoflurane
Ambroxol
Amoxicillin
Fluoruracil
Cisplatin_w
Rivaroxaban
Tobramycin_w
Carboplatin_w
Cyclophosphamide
DMSO+water
Salbutamol_w
Azathioprine
Penicillin G
Vancomycin
Hydrocortisone
Tranexamic acid_w
Piperacillin
Ibuprofen
Moxifloxacin
Domperidone
Imatinib
Acyclovir
Heparin_w
Emtricitabine
Remdesivir
Paclitaxel
Pyrazinamide
Clavulanic acid
DMSO
Ketamine
Oseltamivir
Cephalexin
Abacavir
Acetylsalicylacid
Prednisolone
Dolutegravir
Rocuronium
Glyburide
Rifampicin
Tazobactam
Ascorbic acid
Ramipralat
Bictegravir
Metformin
Nirmatrelvir
Ceftriaxone
Lamivudine
Dapagliflozin
Isoflurane
Metoclopramid
Tenofovir
Empagliflozin
Ciprofloxacin
Dexamethasone
Lisinopril
Caffeine
Molnupiravir
Oxaliplatin_w
Capecitabine
Cefuroxime
Oxymetazoline
Norepinephrine
Dabigatran
Baricitinib
Pantoprazol
Furosemide
Nateglinide
Azithromycin
Losartan
Acarbose
Ritonavir
Metoprolol
Ganciclovir
Clindamycin
Melphalan
Dobutamine
Propranolol
Ampicillin
Mycophenolate
N−Acetyl−L−Cysteine
Acetaminophen
Isoniazid
Metamizole
Omeprazol
Cisatracurium
Epinephrine
Cetirizin
Amethopterin
Propofol
Etomidate
Diclofenac
Fosfomycin_w
Levofloxacin
Enoxaparin_w
Dienogest
Salmeterol
Doxycycline
Montelukast
Loperamide
Tamoxifen
Tacrolimus
Negative control
Bromocriptine
Desogestrel
Doxorubicine
Ethinyl Estradiol
Carvedilol
Levothyroxine
Conc. ANI
0.000 µG/ mL
0.008 µG/ mL
0.020 µG/ mL
0.040 µG/ mL
0.000 µG/ mL control
0.008 µG/ mL control
0.020 µG/ mL control
0.040 µG/ mL control

**Supplementary Figure 3:**

Checkerboard assays with FLC (blue) and ANI (red).

a) Drugs that decreased the fungal biomass with FLC (blue) or ANI (red).

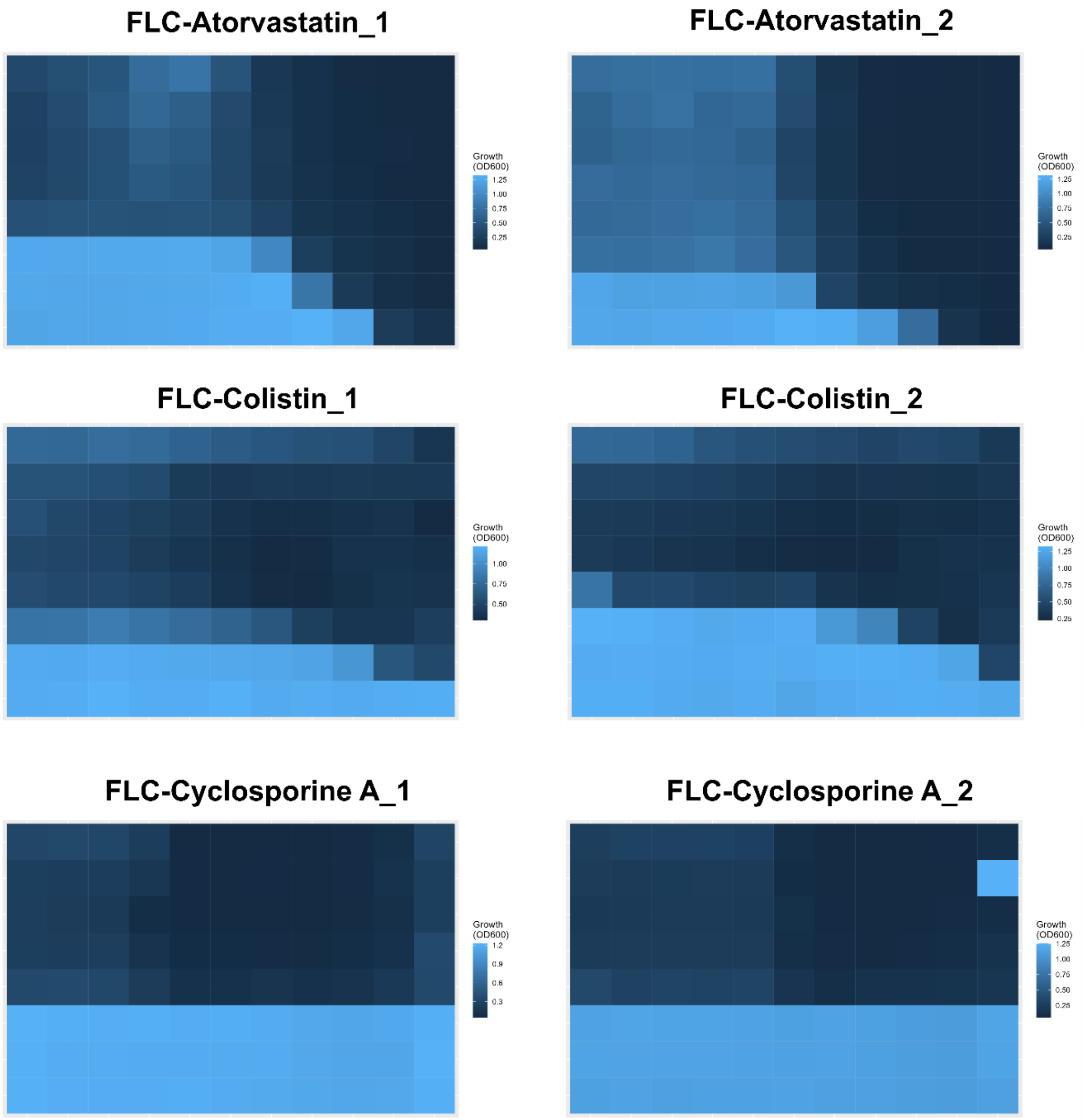

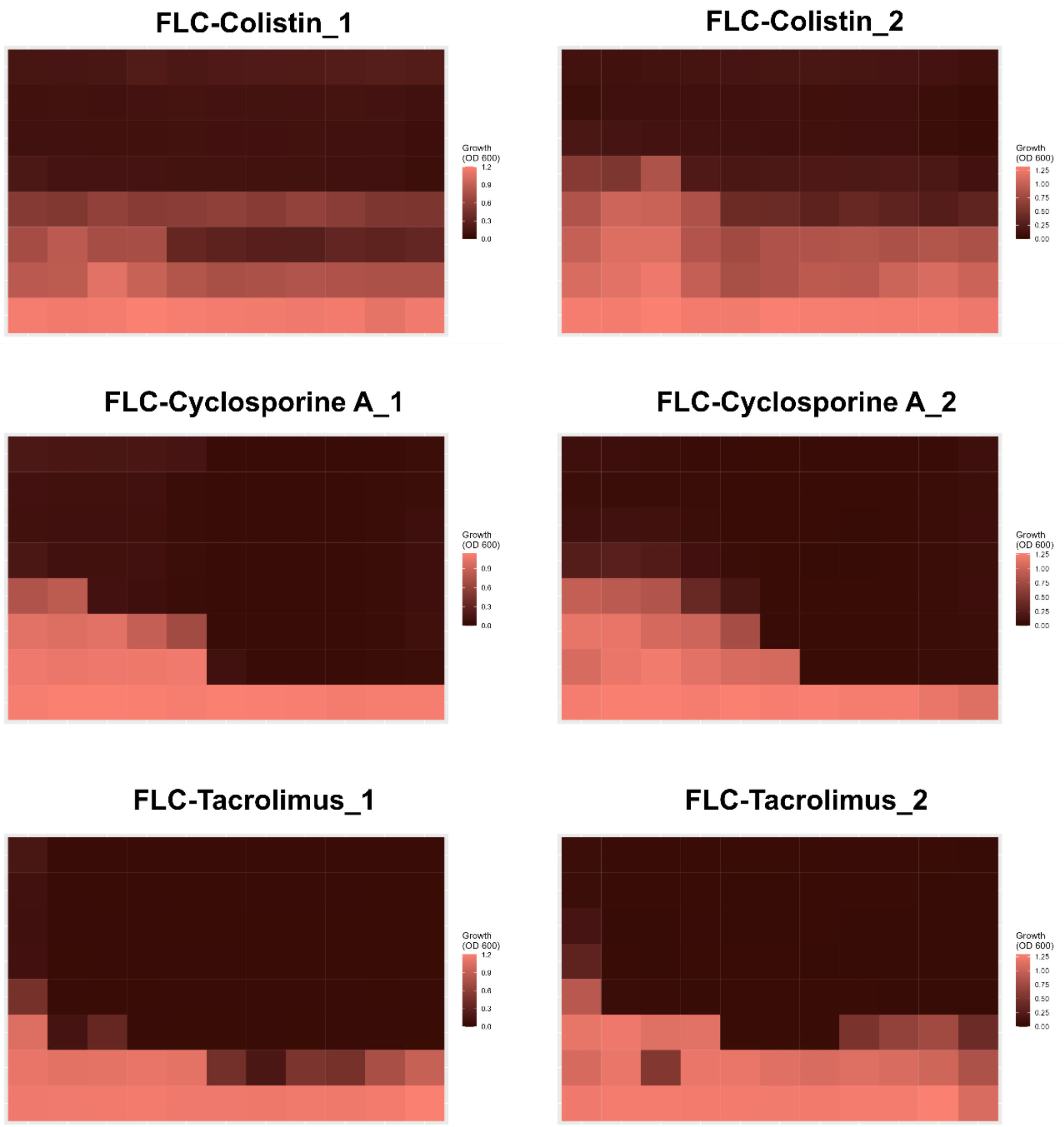
FLC-Colistin_1
FLC-Colistin_2
FLC-Cyclosporine A_1
FLC-Cyclosporine A_2
FLC-Tacrolimus_1
FLC-Tacrolimus_2
Growth (OD 600)
1.2
0.9
0.6
0.3
0.0
1.25
1.00
0.75
0.50
0.25
0.00

b) Durgs that increased the fungal biomass with FLC (blue) or ANI (red).

**FLC-Bromocriptine_1**

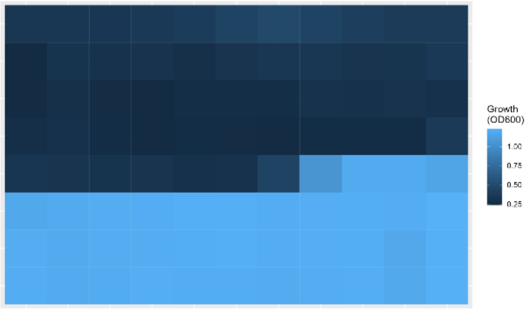


**FLC-Bromocriptine_2**

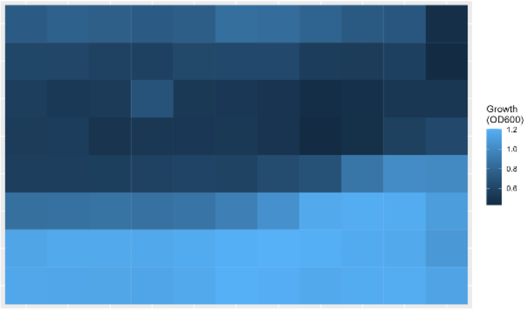


**FLC-Carvedilol_1**

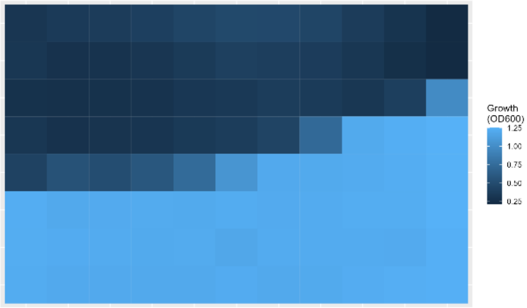


**FLC-Carvedilol_2**

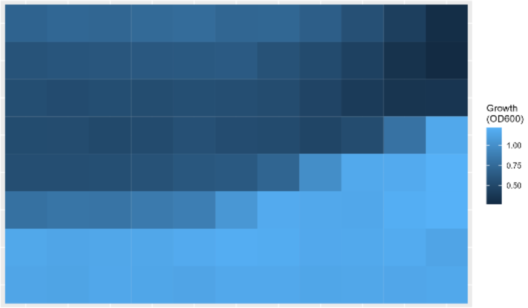


**FLC-Deogestrel_1**

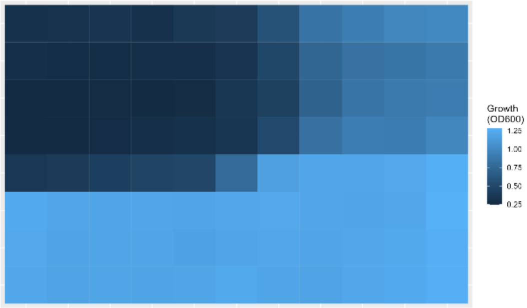


**FLC-Desogestrel_2**

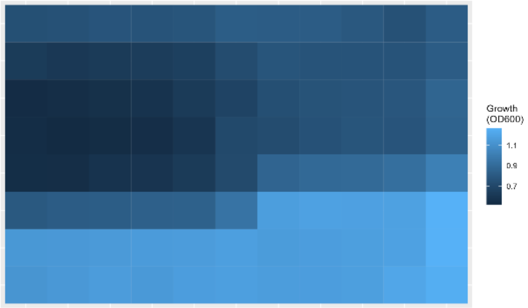


**FLC-Dienogest_1**

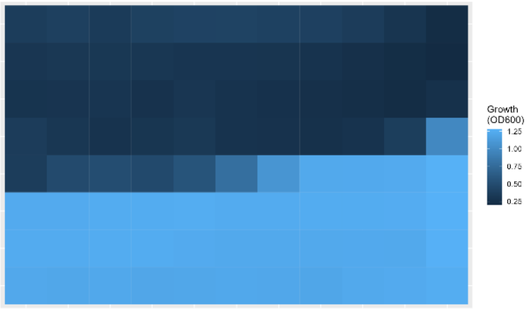


**FLC-Dienogest_2**

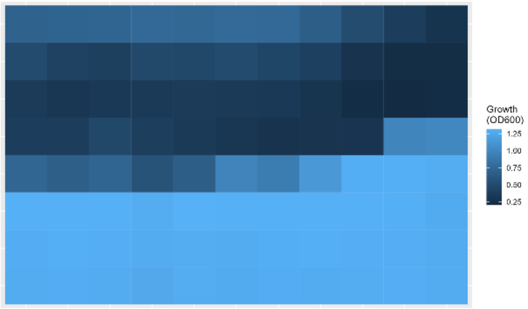

FLC-Doxorubicin_1

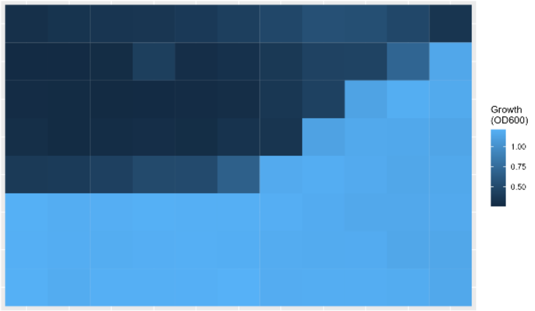


FLC-Doxorubicin_2

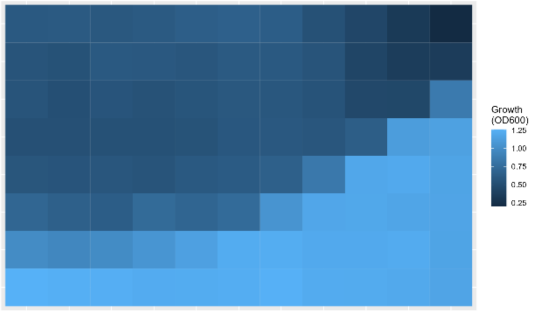


FLC-Doxycycline_1

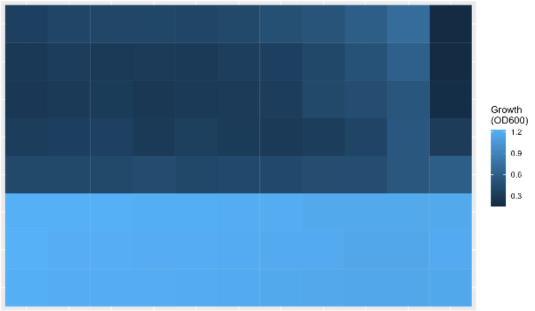


FLC-Doycyline_2

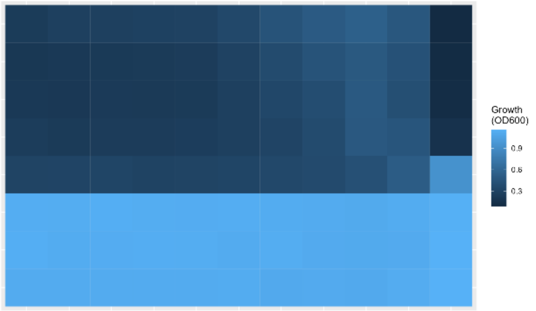


FLC-Enoxaparin_1

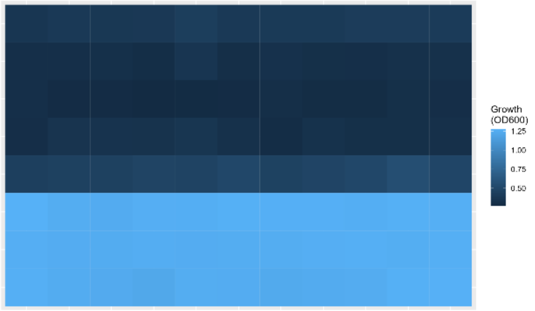


FLC-Enoxaparin_2

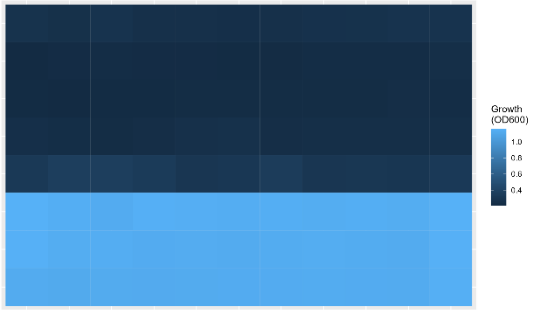


FLC-Estradiol_1

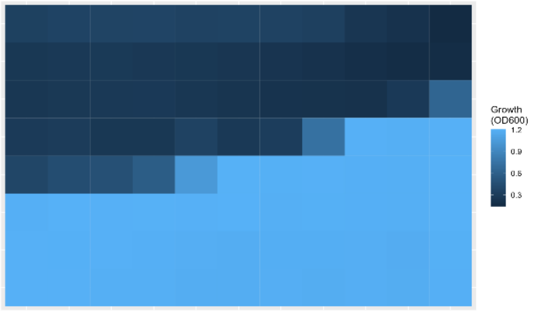


FLC-Estradiol_2

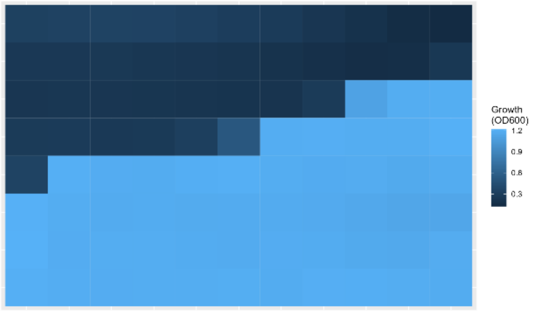

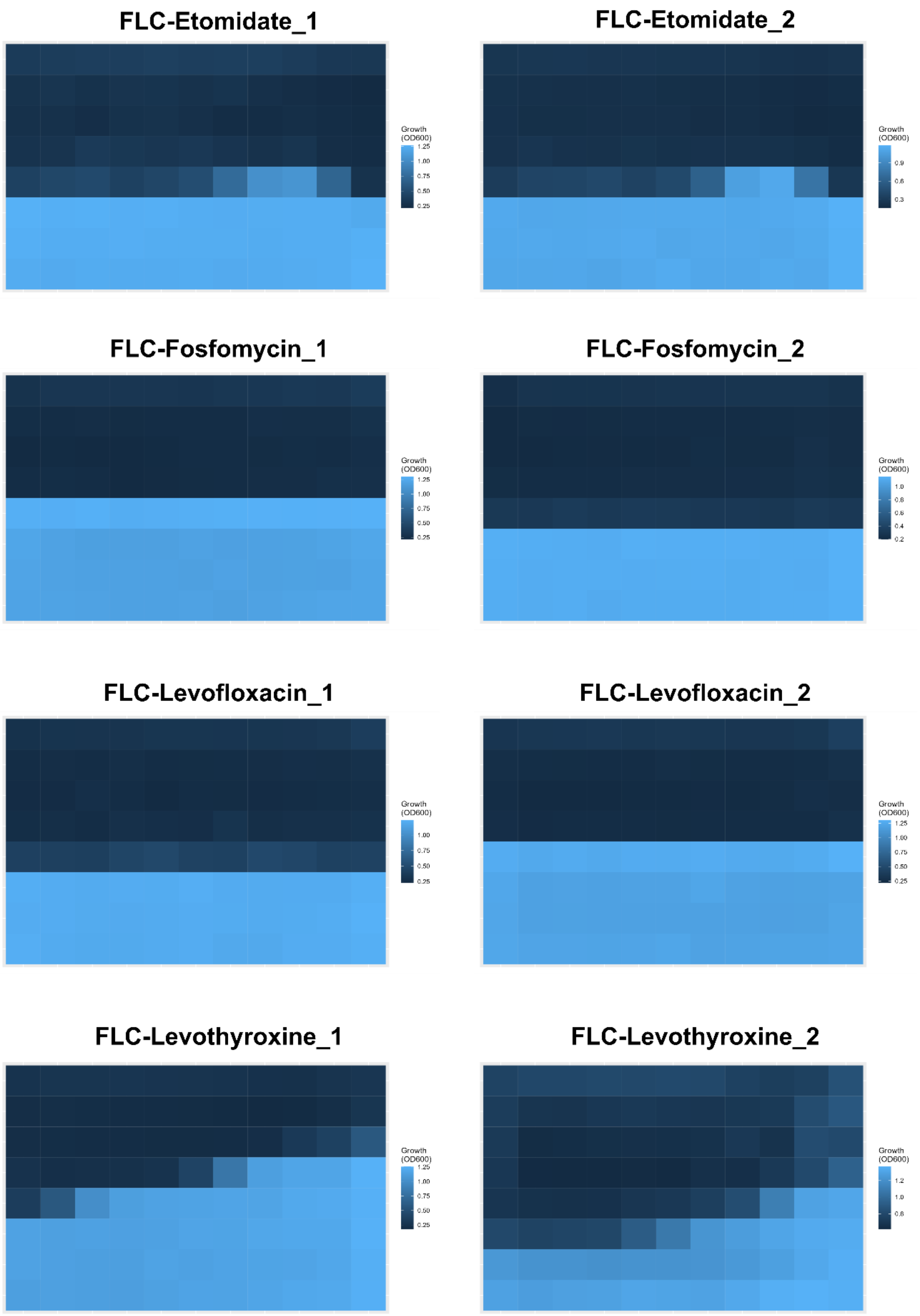
FLC-Etomidate_1
FLC-Etomidate_2
FLC-Fosfomycin_1
FLC-Fosfomycin_2
FLC-Levofloxacin_1
FLC-Levofloxacin_2
FLC-Levothyroxine_1
FLC-Levothyroxine_2
Growth (OD600)

FLC-Loperamide_1

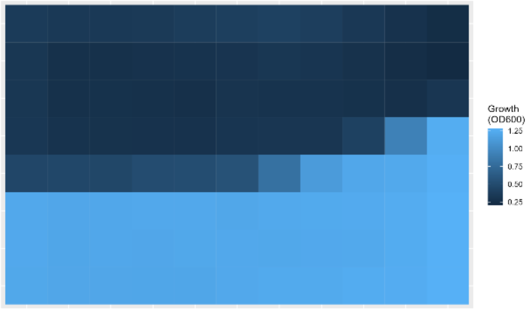


FLC-Loperamide_2

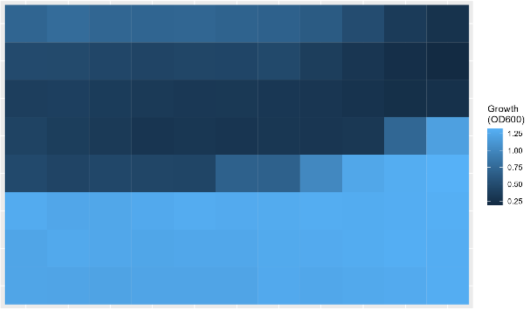


FLC-Montelukast_1

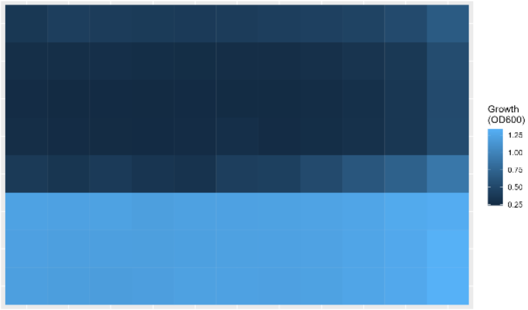


FLC-Montelukast_2

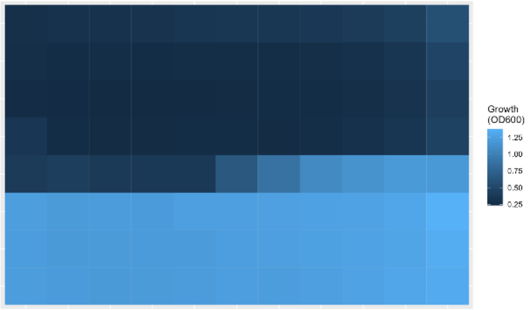


FLC-Mycophenolate_1

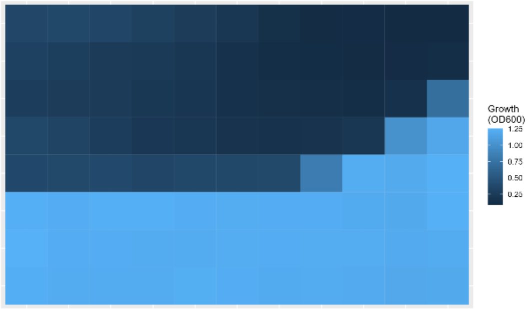


FLC-Mycophenolate_2

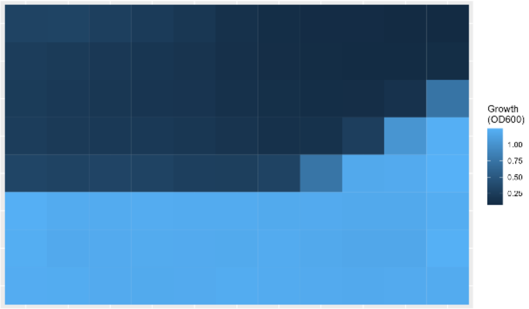


FLC-Salmeterol_1

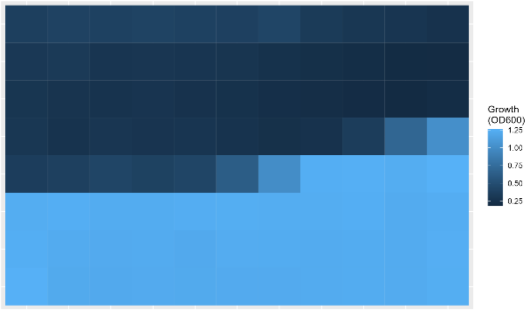


FLC-Salmeterol_2

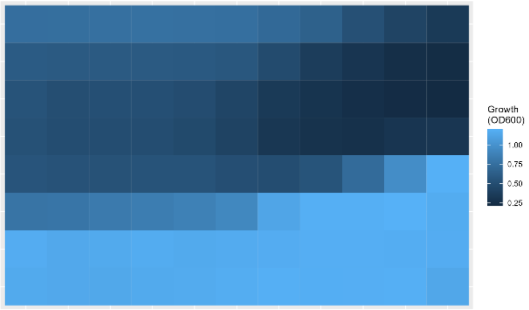

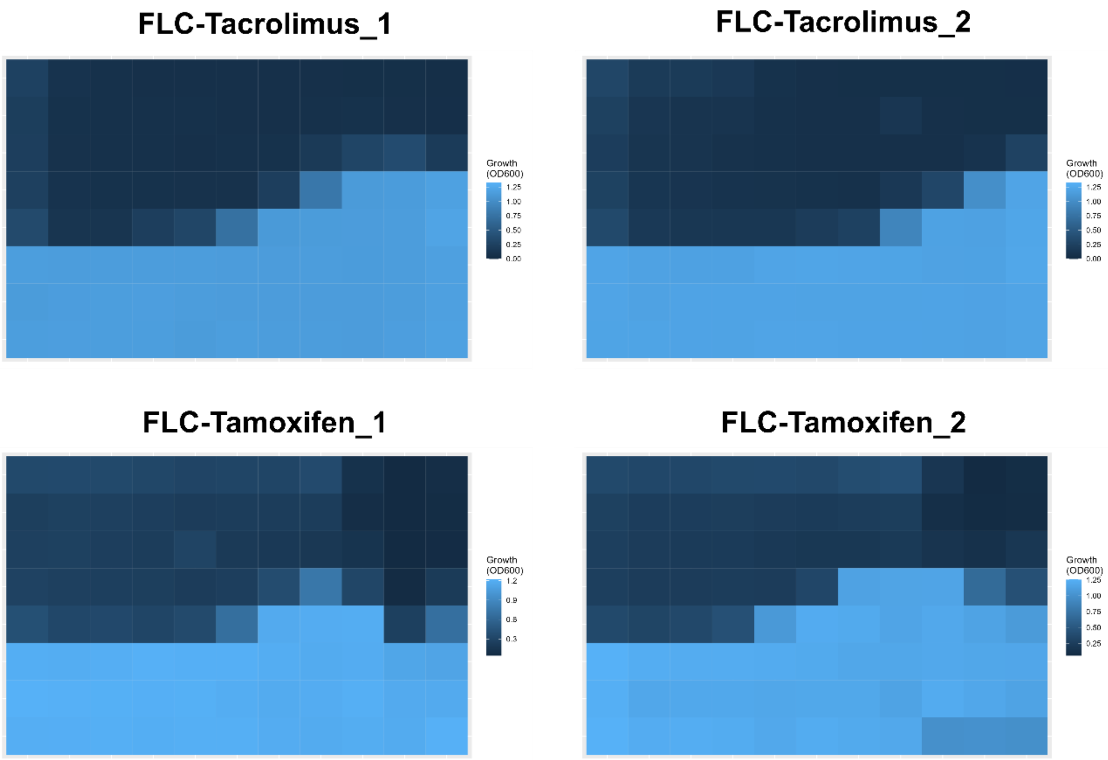
FLC-Tacrolimus_1
FLC-Tacrolimus_2
FLC-Tamoxifen_1
FLC-Tamoxifen_2
Growth (OD600)

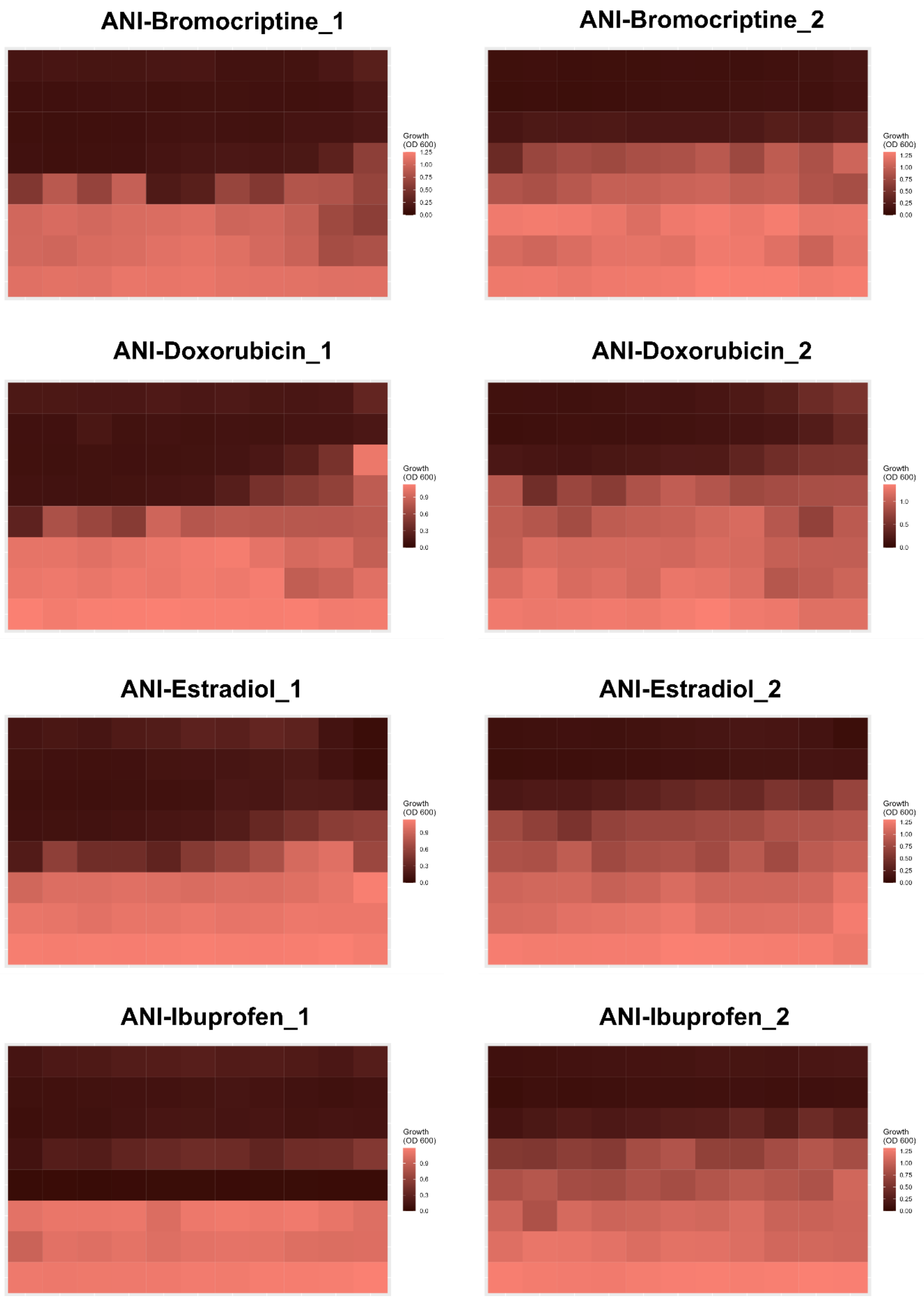
ANI-Bromocriptine_1
Growth (OD 600)
1.25
1.00
0.75
0.50
0.25
0.00
ANI-Bromocriptine_2
Growth (OD 600)
1.25
1.00
0.75
0.50
0.25
0.00
ANI-Doxorubicin_1
Growth (OD 600)
0.9
0.6
0.3
0.0
ANI-Doxorubicin_2
Growth (OD 600)
1.0
0.5
0.0
ANI-Estradiol_1
Growth (OD 600)
0.9
0.6
0.3
0.0
ANI-Estradiol_2
Growth (OD 600)
1.25
1.00
0.75
0.50
0.25
0.00
ANI-Ibuprofen_1
Growth (OD 600)
0.9
0.6
0.3
0.0
ANI-Ibuprofen_2
Growth (OD 600)
1.25
1.00
0.75
0.50
0.25
0.00

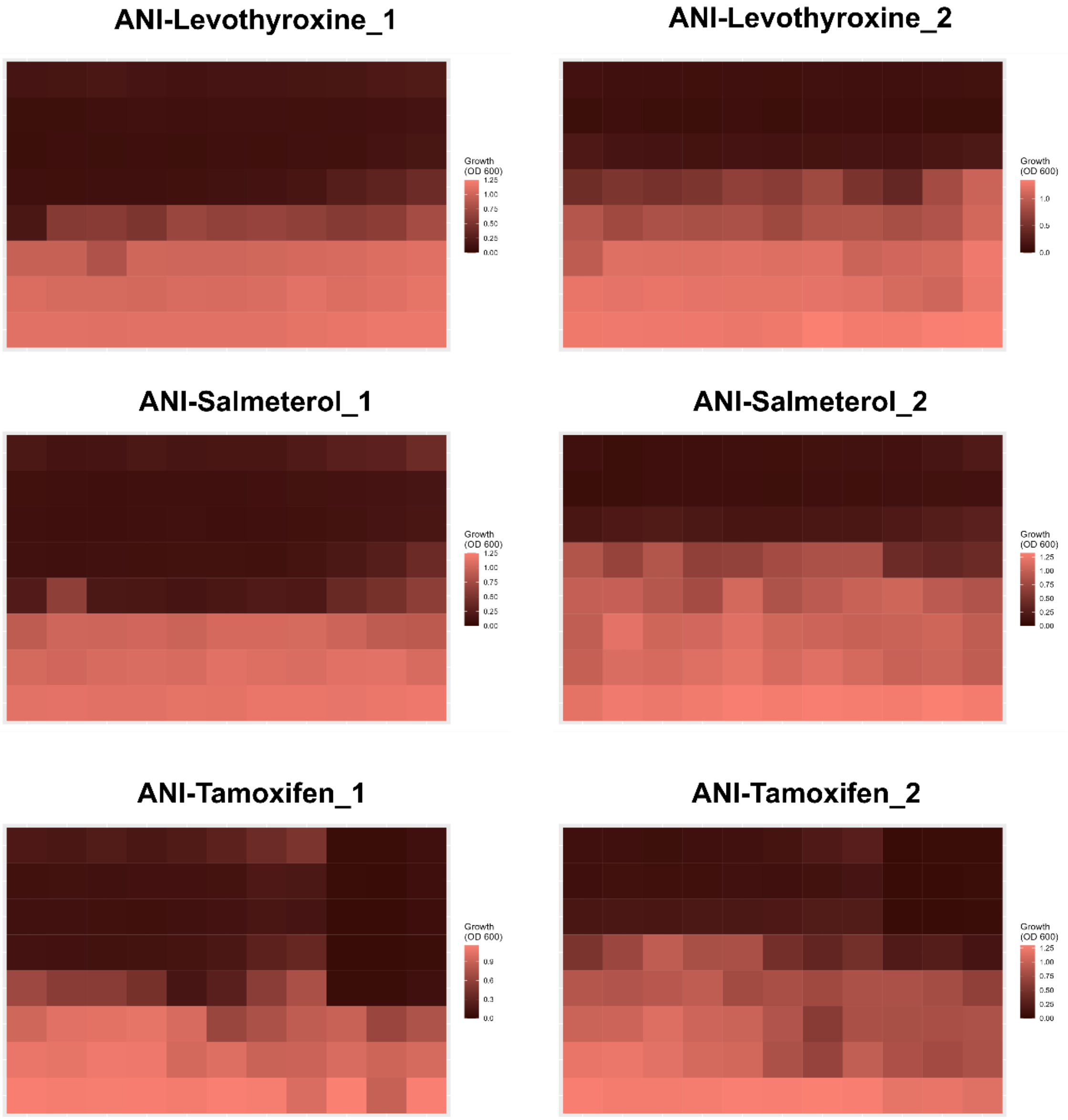
ANI-Levothyroxine_1
ANI-Levothyroxine_2
ANI-Salmeterol_1
ANI-Salmeterol_2
ANI-Tamoxifen_1
ANI-Tamoxifen_2
Growth (OD 600)

**Supplementary Figure 4:**

Growth restoration of negative interactions with FLC (blue) and ANI (red) compared to the untreated control.

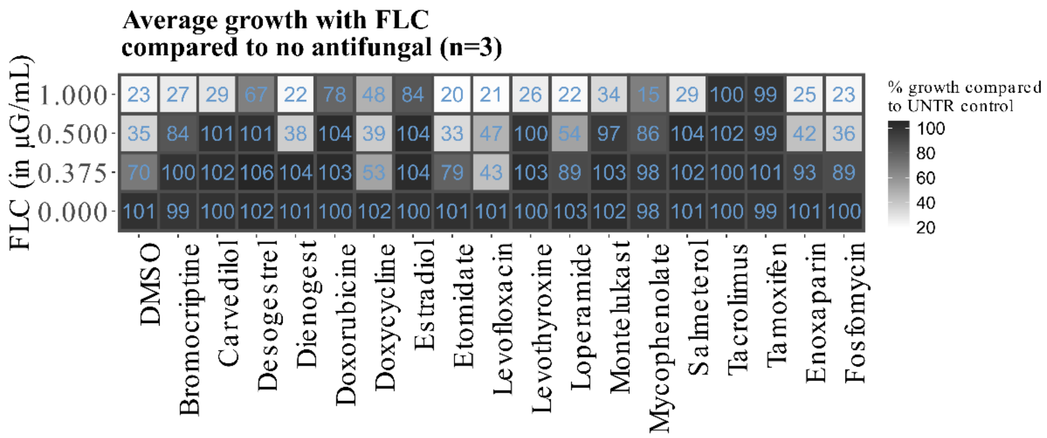


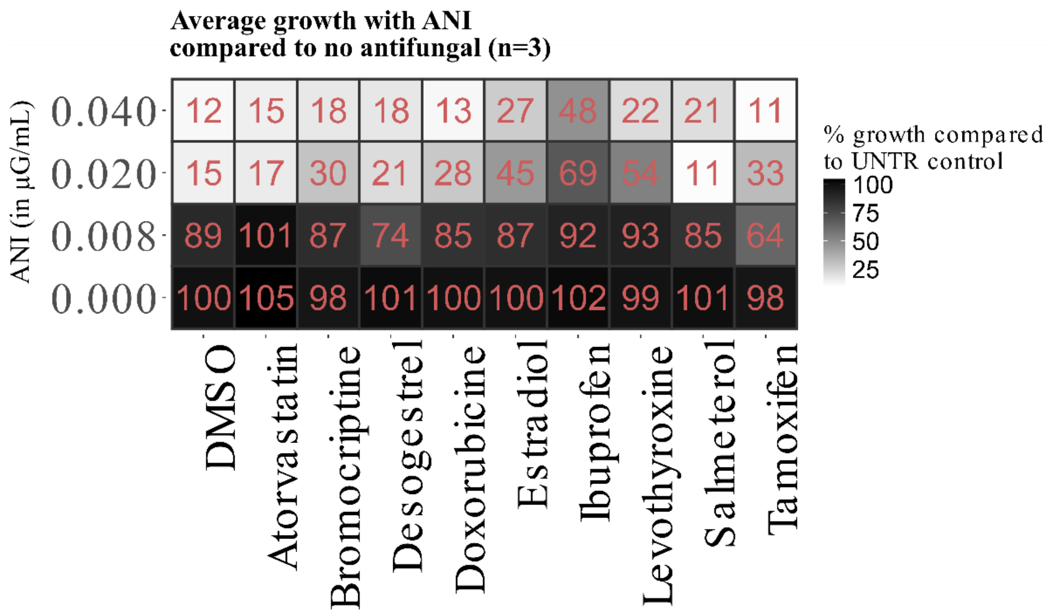

**Supplementary Figure 5:**

Disk diffusion assays with FLC and ANI.

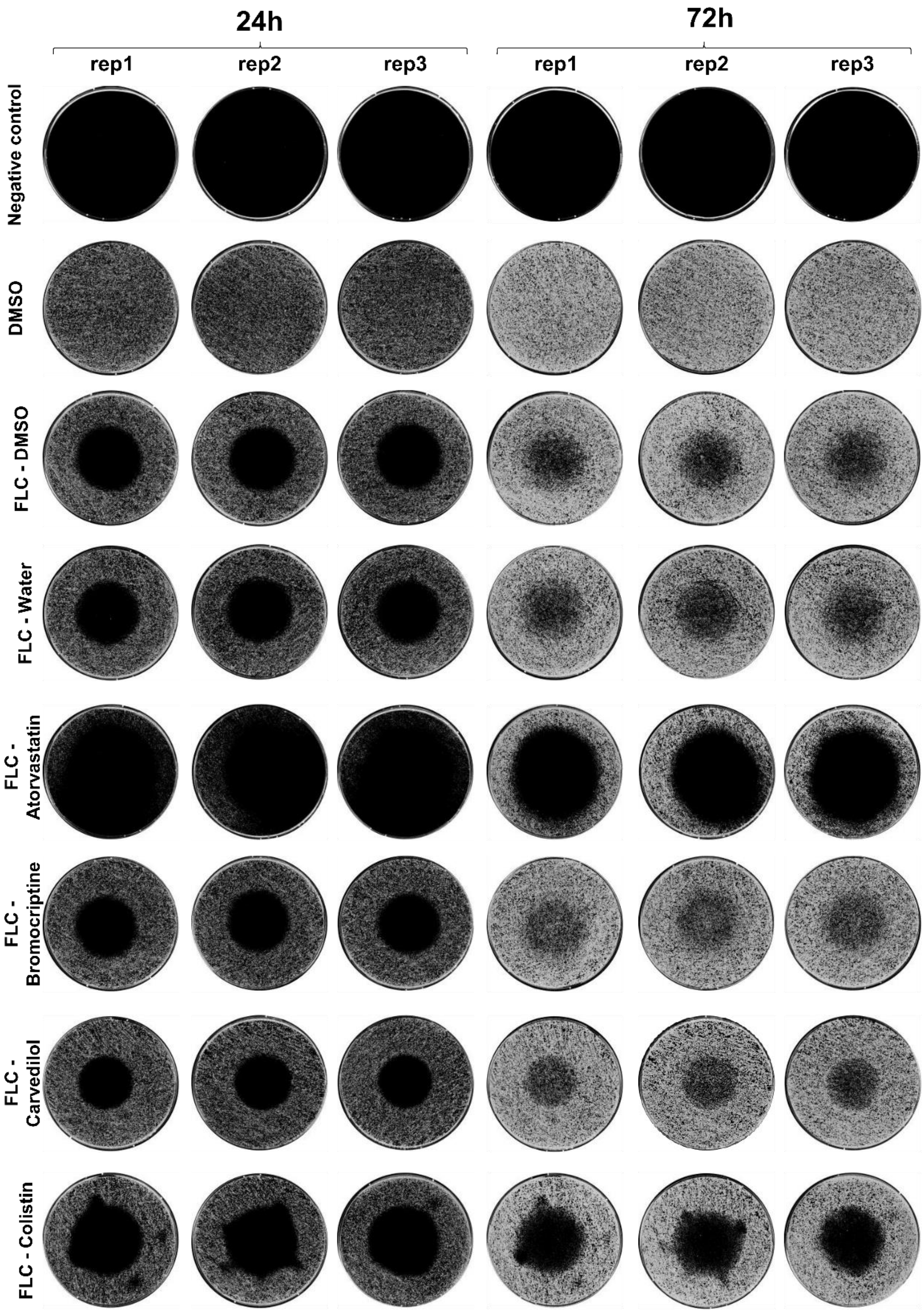

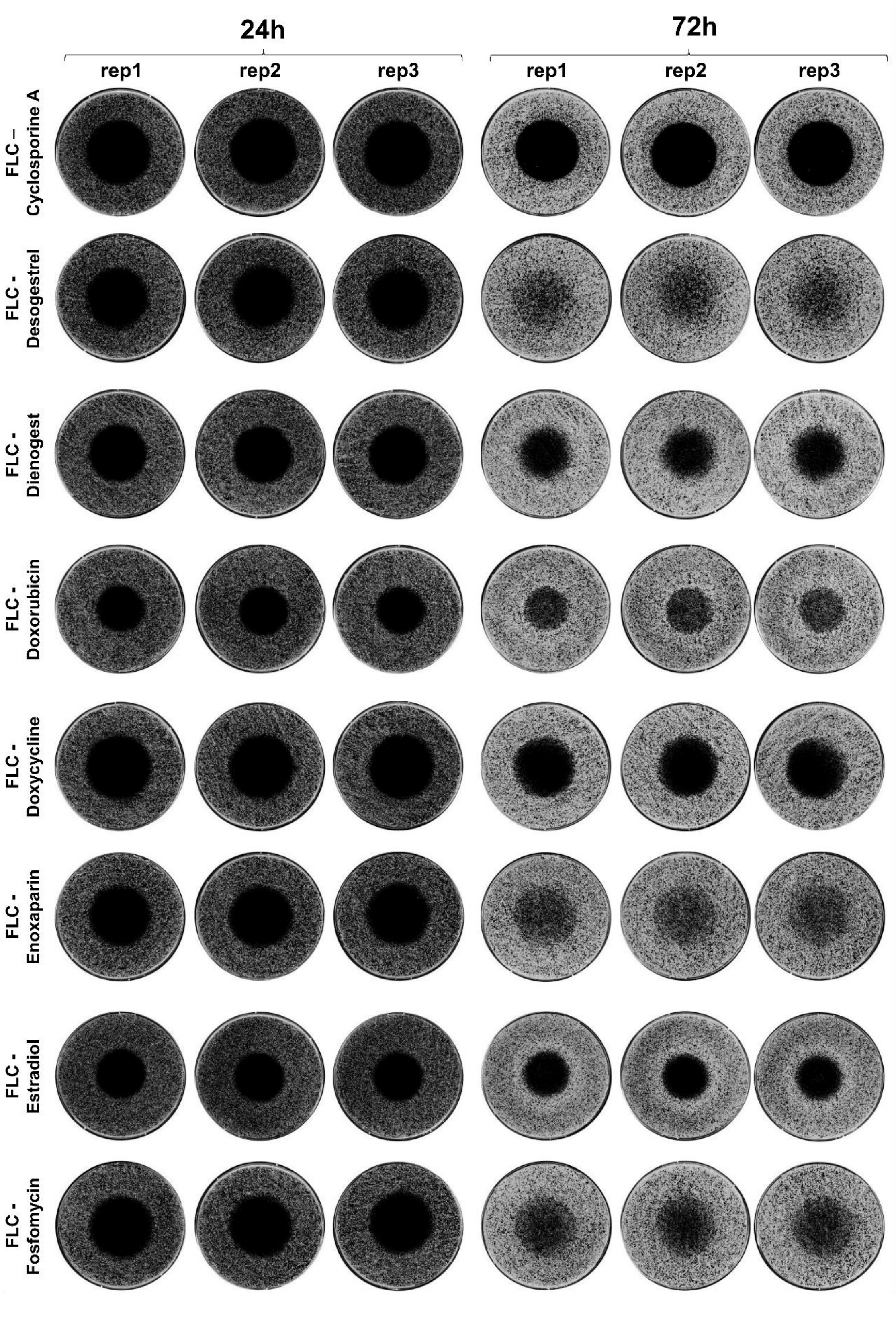
24h
72h
rep1
rep2
rep3
rep1
rep2
rep3
FLC – Cyclosporine A
FLC - Desogestrel
FLC - Dienogest
FLC - Doxorubicin
FLC - Doxycycline
FLC - Enoxaparin
FLC - Estradiol
FLC - Fosfomycin

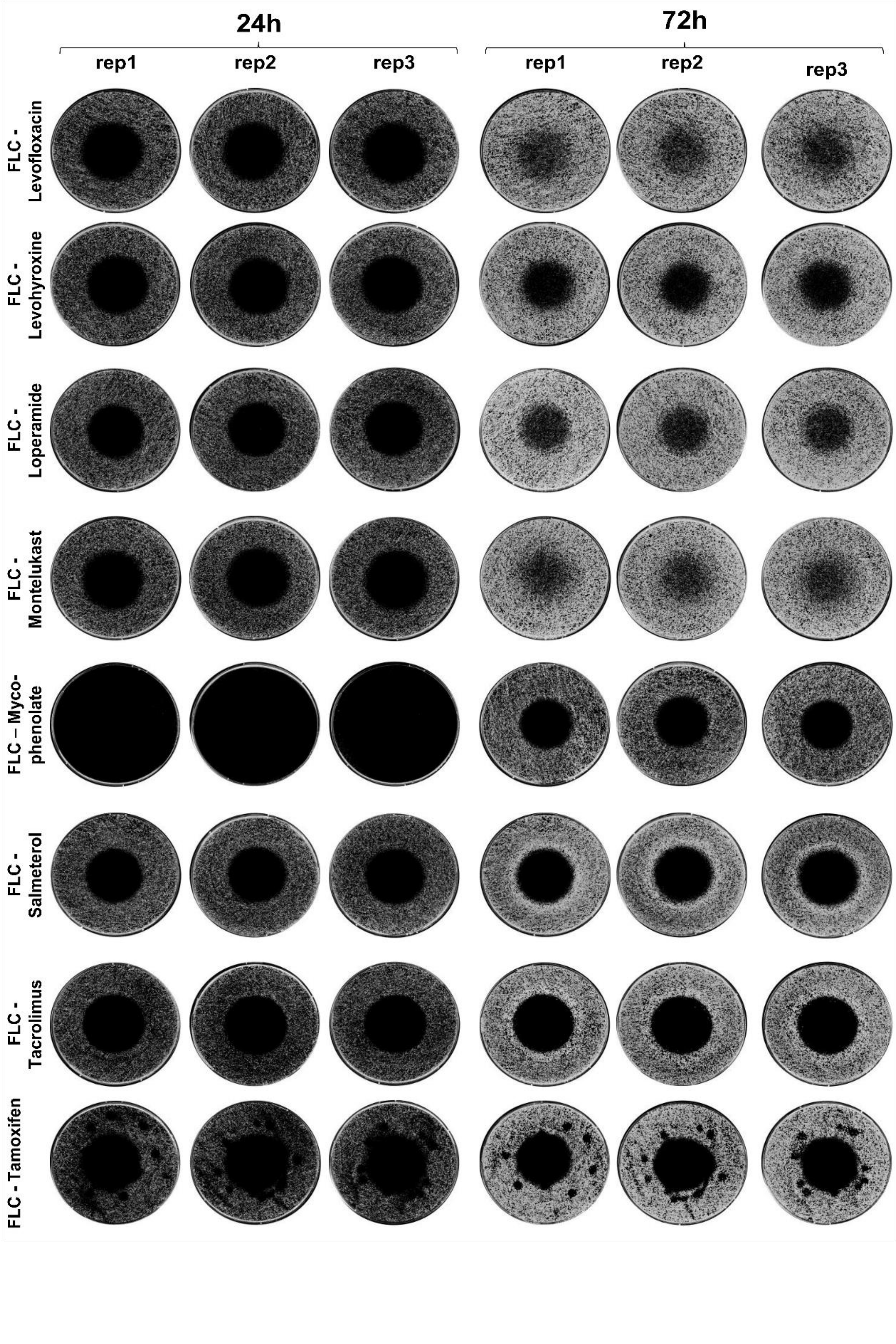
24h
72h
rep1
rep2
rep3
rep1
rep2
rep3
FLC - Levofloxacin
FLC - Levohyroxine
FLC - Loperamide
FLC - Montelukast
FLC – Myco-phenolate
FLC - Salmeterol
FLC - Tacrolimus
FLC - Tamoxifen

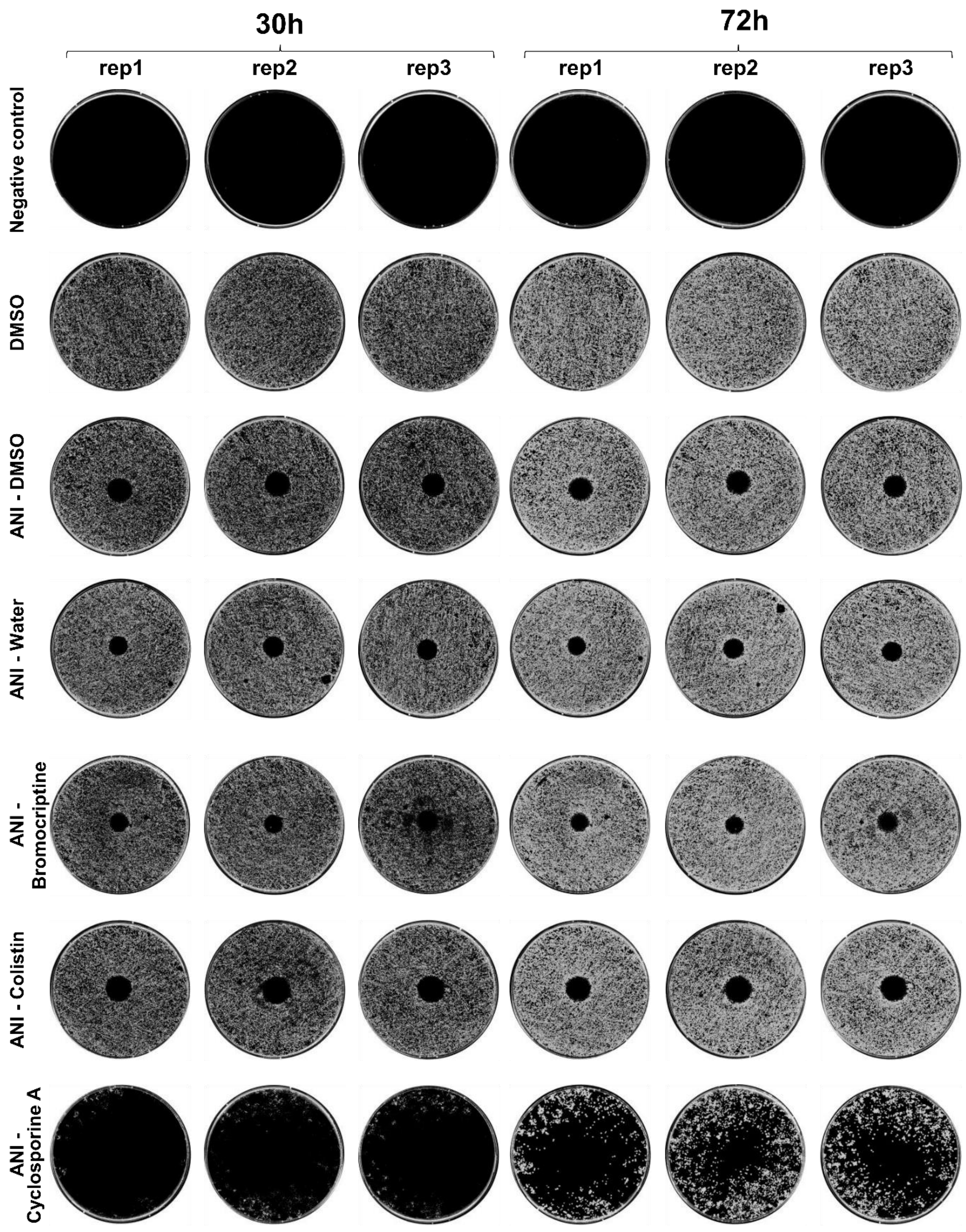
30h
72h
rep1
rep2
rep3
rep1
rep2
rep3
Negative control
DMSO
ANI - DMSO
ANI - Water
ANI - Bromocriptine
ANI - Colistin
ANI - Cyclosporine A

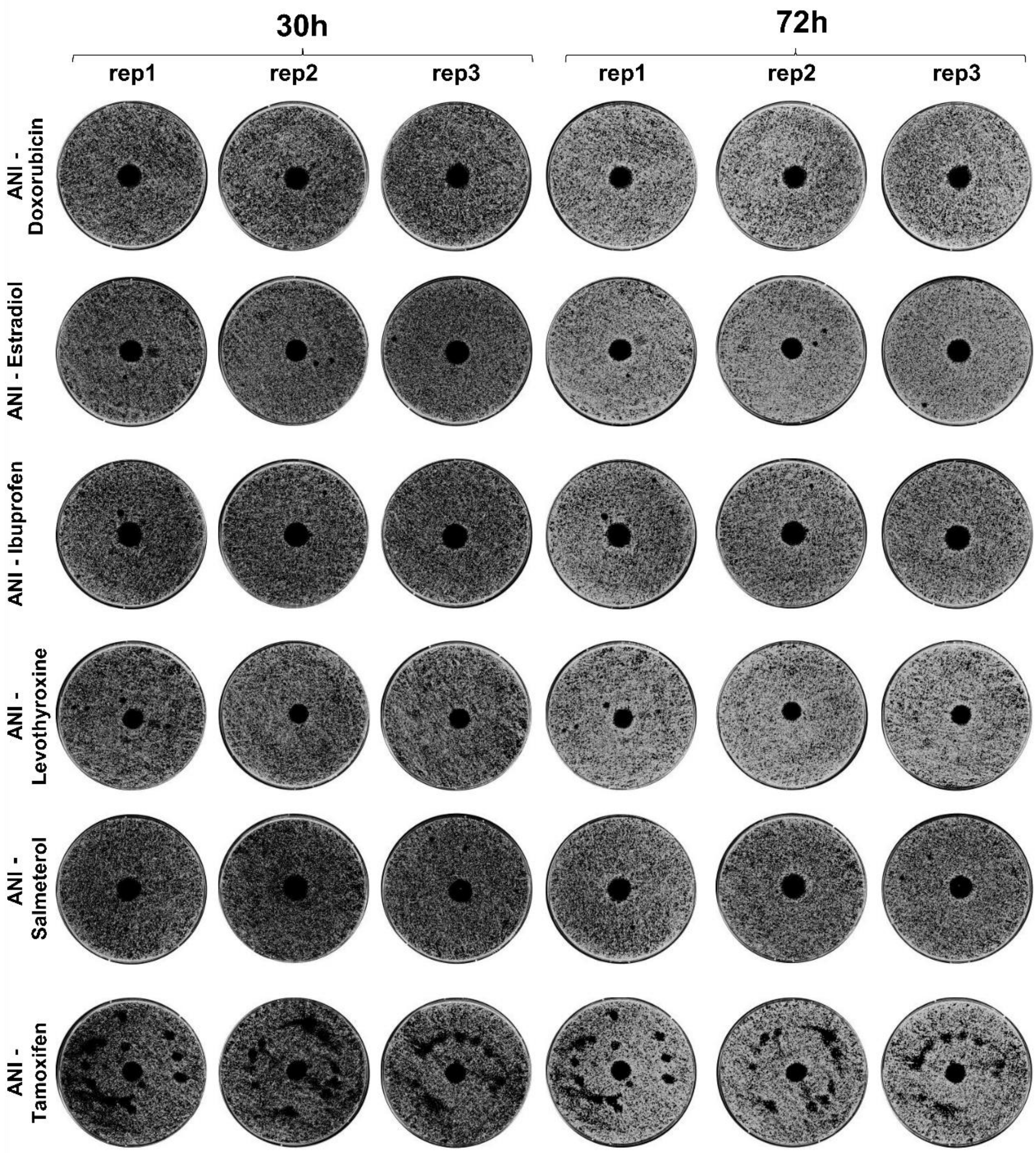
30h
72h
rep1
rep2
rep3
rep1
rep2
rep3
ANI - Doxorubicin
ANI - Estradiol
ANI - Ibuprofen
ANI - Levothyroxine
ANI - Salmeterol
ANI - Tamoxifen

**Supplementary Figure 6:**

In-vivo tests of FLC antagonists in *G. mellonella* larvae. *G. mellonella* were infected with *C. albicans* and treated with FLC alone, FLC + FLC antagonist, FLC antagonist only or not treated. **(A)** Time of survival up to 50% death of larvae show that most proposed FLC antagonists also shortened the time of *G. mellonella* median survival in most drugs. **(B)** Kaplan-Meier survival analysis of different tested drugs. Compounds which differed significantly from the FLC treatment only are shown in Figure 5. A Wilcoxon rank sum test and a Bonferroni-correction was performed to test statistical differences between the groups, indicated as different letters above the graphs. N=15, 2 replicates per experiment. Right-censored data are depicted as crosses.

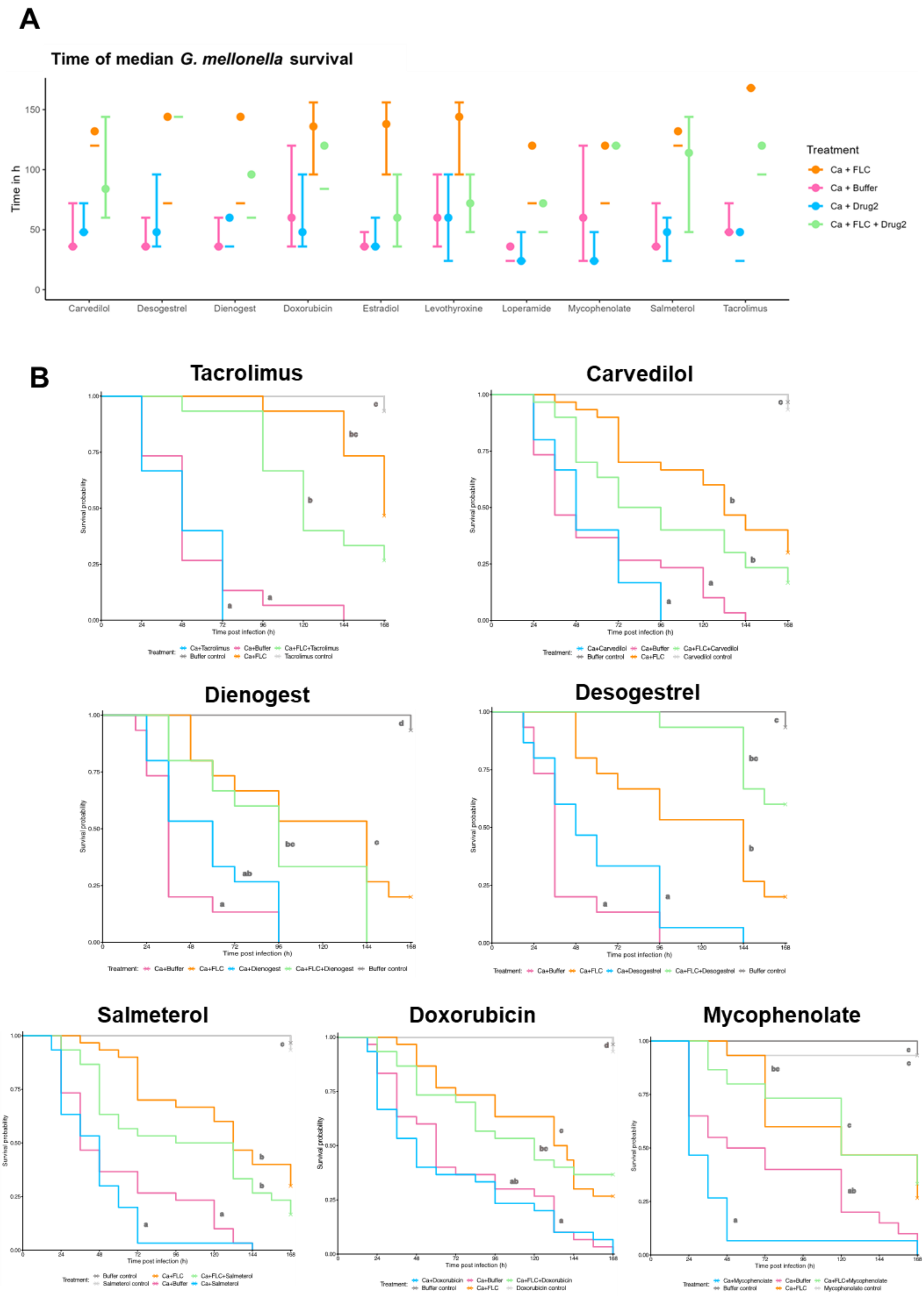
A
Time of median G. mellonella survival
Time in h
Carvedilol
Desogestrel
Dienogest
Doxorubicin
Estradiol
Levothyroxine
Loperamide
Mycophenolate
Salmeterol
Tacrolimus
Treatment
Ca + FLC
Ca + Buffer
Ca + Drug2
Ca + FLC + Drug2
B
Tacrolimus
Carvedilol
Dienogest
Desogestrel
Salmeterol
Doxorubicin
Mycophenolate
Survival probability
Time post infection (h)
